# Fast, Bright and Reversible Rhodamine Tags for Live-Cell Imaging

**DOI:** 10.1101/2025.07.06.663254

**Authors:** Julian Kompa, Lars Jeremy Dornfeld, Nicola Porzberg, Soohyen Jang, Simon Hans Lilje, Claudia Catapano, David Jocher, Lukas Merk, Silja Zedlitz, Runyu Mao, Jonas Wilhelm, Marina Dietz, Miroslaw Tarnawski, Julien Hiblot, Mike Heilemann, Kai Johnsson

## Abstract

We present Rho-tag and SiR-tag, engineered protein tags derived from bacterial multidrug-resistance proteins that bind unsubstituted (silicon-) rhodamines with nanomolar affinity, enabling fast, reversible, and fluorogenic protein labeling. In live cells, Rho-tag labeling occurs within seconds — faster than HaloTag7 — and the tags are compatible with super-resolution methods like STED, SMLM, and MINFLUX. The high specificity of Rho-tag and SiR-tag for unsubstituted rhodamines allows their use alongside HaloTag7 and SNAP-tag. *In vivo* applications are demonstrated by efficient neuronal labeling in zebrafish larvae.

## Introduction

The development of fluorescent probes and labeling strategies has been fundamental to advances in bioimaging.^[1]^ Fluorescent proteins (FPs) remain the most popular fluorescent probes as they can be genetically targeted to the protein or cell of interest.^[2]^ Synthetic fluorophores, particularly those in the red and far-red spectral regions, are brighter and more photostable than FPs, but require a labeling strategy.^[3]^ Amongst the chemical dyes, rhodamine probes stand out due to their excellent brightness, cell-permeability and broad spectral range.^[4]^ Another key feature is that they exist in an environmentally-sensitive equilibrium between a fluorescent zwitterion (open) and a non-fluorescent (closed) yet cell-permeable spirolacton form.^[5]^ This has been extensively explored for the generation of fluorogenic rhodamine derivatives.^[6, 7]^ Rhodamine probes are typically targeted using self-labeling protein (SLP) tags like HaloTag^[8]^ or SNAP-tag^[9]^. While these tags enable relatively fast and efficient covalent labeling of fusion proteins, they require the derivatization of rhodamines with a substrate for the SLPs such as the HaloTag Ligand (HTL). As a consequence, the resulting rhodamine derivatives are larger molecules with a number of features that make labeling *in vivo* relatively inefficient. An alternative to the labeling of SLPs with rhodamines is the use of proteins that directly bind underivatized fluorophores with high binding affinity like fluorogen-activating proteins (FAPs).^[10]^ A prominent example is the fluorescence-activating and absorption-shifting tag (FAST), which allows the targeting of FP-like fluorogens of different colors using a single protein tag.^[11]^ However, the spectroscopic properties of the underlying fluorophores are inferior to those of rhodamines.

Here, we present two rhodamine-binding protein-tags (Rho-tag and SiR-tag) for live-cell and *in vivo* protein labeling with tetramethyl rhodamine (TMR) and silicon rhodamine (SiR). Rho-tag and SiR-tag bind their substrates with high affinity, specificity and speed. Binding of TMR and SiR by their respective tags improves the brightness of the fluorophores, and Rho-tag and SiR-tag can be used in super-resolution microscopy, including single-molecule localization microscopy (SMLM)^[12]^, stimulated emission depletion (STED) microscopy^[13]^ and MINFLUX^[14]^. Due to their fast labeling and good permeability, TMR and SiR efficiently label the two tags in neurons of zebrafish larvae. Rho-tag and SiR-tag thus open up new applications for rhodamines in bioimaging.

## Results

### Candidate selection and Rho-tag engineering

*In silico* pharmacokinetic analysis of TMR-HTL (in its open form) and SiR-HTL (in its closed form, Figure 1A) using the SwissADME^[15]^ webtool confirmed the experimentally observed relatively poor brain-penetration of these molecules (Figure 1B, Table S1). However, the unsubstituted fluorophores without any HaloTag Ligand (HTL, Figure 1A) were predicted to readily cross the blood-brain barrier. This suggests that a protein with high affinity for unsubstituted rhodamines, like TMR or SiR, could be advantageous for labeling applications *in vivo*.

**Figure 1.**
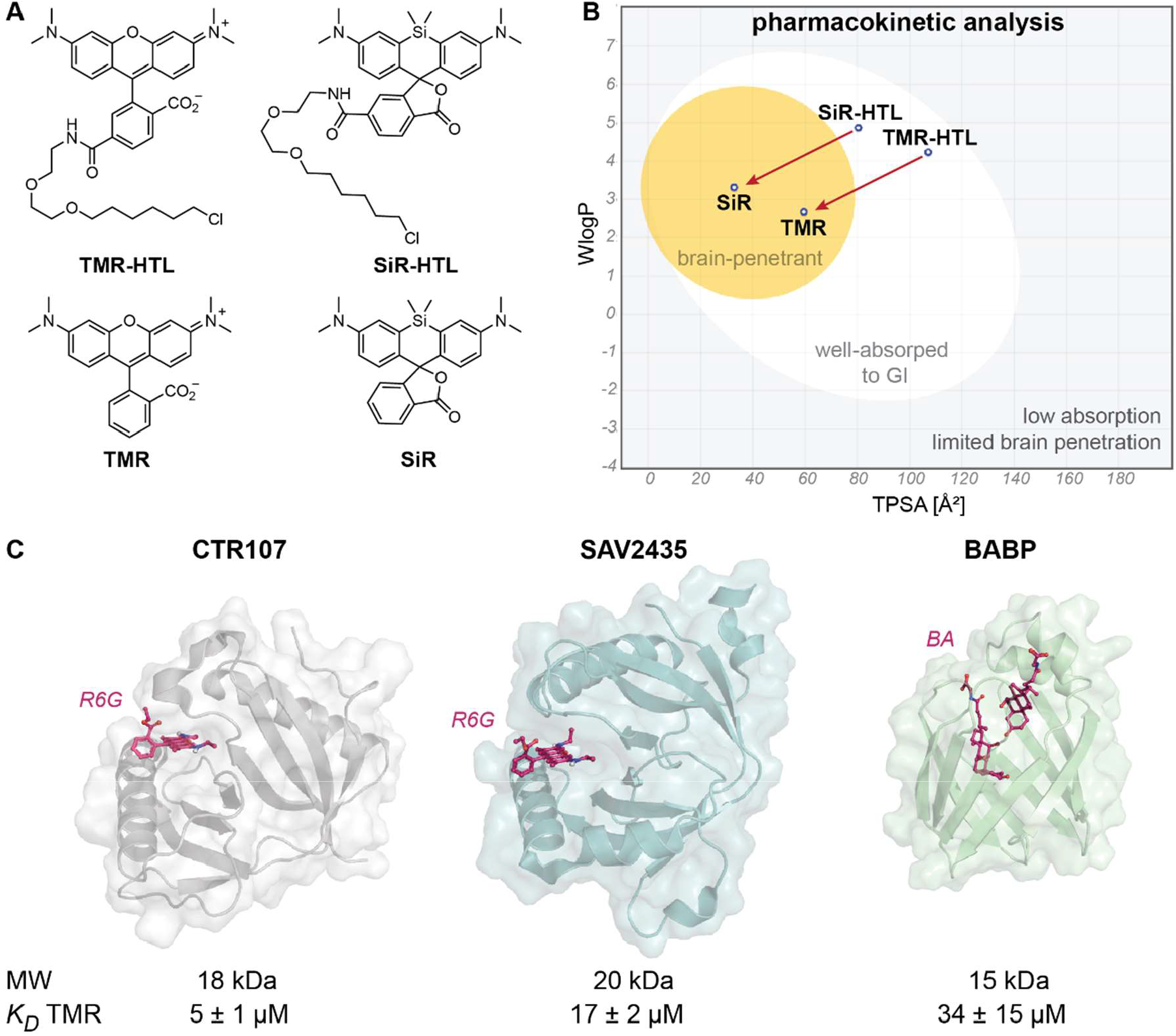
Dye and candidate selection for the development of Rho-tags. A) Chemical structure of TMR and SiR and their corresponding HaloTag Ligand (HTL) derivatives. SiR is shown in its closed form (spirolacton), whereas TMR is shown in its open, zwitterionic form. B) BOILED-Egg^[40]^ representation generated with SwissADME^[15]^ for molecules shown in (A). Correlation between topological polarity (TPSA) and predicted lipophilicity (WlogP). Gray area – predicted low passive absorption to the gastrointestinal (GI) system and limited brain access (BBB). Egg white - high probability for GI absorption. Egg yolk – high probability for BBB permeation. Blue dot – substrate is predicted to be actively exported from the central nervous system. C) Ligand-bound structures, size (MW) and binding affinity to TMR (*K*_*D*_) of three potential Rho-tag candidates. Two multidrug binding proteins CTR107 (PDB-ID 5KAX, X-ray, 2.0 Å resolution) and SAV2435 (PDB-ID 5KAW, X-Ray, 1.8 Å resolution) in complex with Rhodamine 6G (R6G). Bile-acid binding protein (BABP, PDB-ID 2LFO, NMR) in complex with bile acids (glycochenodeoxycholic and glycocholic acids, BA). Tertiary structures are represented as colored cartoons and surface. The ligands are represented as purple sticks. The binding affinity *K*_*D*_ was derived from fluorescence polarization experiments at constant dye concentration (20 nM) and serial protein dilutions (0 – 200 µM). Average data and standard deviation from at least three independent measurements (N ≥ 3). CTR107 was selected as lead candidate for the development of Rho-tags due its moderate binding affinity, small size and single domain state.

In our search for a suitable starting point for generating a so-called Rho-tag, we identified three proteins known to bind rhodamines from literature resources and the protein data bank (PDB)^[16]^. CTR107 and SAV2435 are multi-drug resistant (MDR) proteins capable of polyspecific substrate binding through their Gyrl-like domains.^[17]^ Both proteins display a globular fold of 18 kDa and 20 kDa, respectively, and have been co-crystalized with rhodamine 6G (R6G, Figure 1C).^[17]^ The third protein, bile-acid binding protein (BABP), has been exploited to encapsulate rhodamine dyes on functionalized surfaces (Figure 1C).^[18]^ These three proteins were recombinantly expressed and evaluated for their TMR binding. SAV2435 and BABP showed weak binding to TMR (*K*_*D*_ 17 ± 2 µM and 34 ± 15 µM, respectively). In addition, SAV2435 quenched the TMR fluorescence, presumably through a tryptophan residue in the binding site (Figure S1). CTR107 displayed a *K*_*D*_ of 5.2 ± 0.7 µM for TMR, which makes it a promising scaffold for the development of a Rho-tag. We tested the CTR107 binding affinity for a panel of standard dyes and found that it binds to different rhodamines such as rhodamine B and R101, and pyronines (Figure S2). However, compared to TMR and SiR, these dyes displayed strong cellular background when incubated with U2OS cells, restricting their use for live-cell fluorescence imaging (Figure S3). We therefore focused on the generation of high-affinity TMR and SiR binders.

Next, we performed a structure-guided site-directed mutagenesis screening on CTR107 (Figure S4), and identified mutations E36V and N138A which improved the TMR binding by 48-fold (*K*_*D*_ 108 ± 29 nM). This first-generation Rho-tag0.1 was expressed in live U2OS cells as a histone2B (H2B) fusion and could be specifically labeled with TMR, which was not possible with the parental CTR107 (Figure S5). As we observed minor background staining from unbound fluorophore, we attempted to further increase the binding affinity of Rho-tag0.1 by directed evolution (Figure S6). This would allow to label at lower TMR concentrations. Two types of Rho-tag0.1 libraries were displayed on the surface of yeast and subjected to iterative rounds of fluorescence-activated cell sorting (FACS, supplementary note 1). First, we generated site-saturation mutagenesis libraries (SSMLs) targeting the 16 amino acid alpha-helix *Hα1* (^31^LGSLFVAGYHDILQLL^46^) near the fluorophore binding site. From these libraries we identified the double mutation S33N / H40T which further improved the binding affinity towards TMR by 6-fold (*K*_*D*_ 19 ± 9 nM). Second, we screened a synthetic deep mutational scanning (DMS) library covering the full protein, from which we identified the M64F mutation. To offset a decrease in thermostability observed throughout the engineering process, we employed the Protein Repair One-Stop Shop (PROSS)^[19]^, to generate protein variants with increased thermostability but retaining similar binding affinities (supplementary note 2, Table S2). Combining the previously discovered E36V / N138A with S33N / H40T / M64F mutants and 14 additional mutations suggested by the PROSS designs yielded the final Rho-tag having a melting temperature of 61 °C.

### Rho-tag is a bright and specific binder to TMR for cellular labeling

Rho-tag features 19 point-mutations compared to the parental protein (Figure 2A). The *N*-terminal cysteine residues (position 5 and 9) can be mutated to serine without impairing the binding function (supplementary note 3). To understand the molecular basis of the ligand binding, we solved the crystal structure of Rho-tag in complex with TMR (Figure 2B, PDB ID 9RTM, 2.1 Å resolution) which reveals deep insertion of the xanthene into the binding pocket (Figure 2C). We found a putative hydrogen bond interaction of the α-amino group of A140 with the *ortho*-carboxy moiety of TMR and hypothesize, that this interaction is important to stabilize the dye in its open, fluorescent form (Figure 2D). Rho-tag possesses a *K*_*D*_ of 2.7 ± 2.1 nM for TMR, a ∼2000-fold improvement relative to parental CTR107. We further investigated the binding kinetics of TMR *in vitro*. Stopped-flow fluorescence anisotropy experiments revealed on-rates close to the diffusion limit (*k*_*on*_ 1.4 ± 0.5 × 10^8^ M^-1^s^-1^). The off-rate *k*_*off*_ of TMR was calculated from the *K*_*D*_ and the *k*_*on*_ as 0.4 ± 0.3 s^-1^ (Table 1). The quantum yield of TMR increased upon binding to Rho-tag from 0.44 ± 0.01 to 0.73 ± 0.02, resulting in an almost two-fold increase in brightness (Table 1). This makes Rho-tag about 36% brighter than TMR-labeled HaloTag7 *in vitro*. Rho-tag can also be labeled with JF_549_^[20]^, which shows similar binding affinity as TMR (*K*_*D*_ 4.0 ± 1.2 nM). In order to gain greater insights into the protein-ligand interactions, we performed classical molecular dynamics (MD) simulations using the Rho-tag / TMR complex crystal structure. The ligand RMSD (root mean square deviation) remains relatively constant (∼0.3 Å) throughout the simulation. All rotatable bonds are conformationally restrained (Figure S7), providing a rational for the high quantum yield.

**Table 1.**
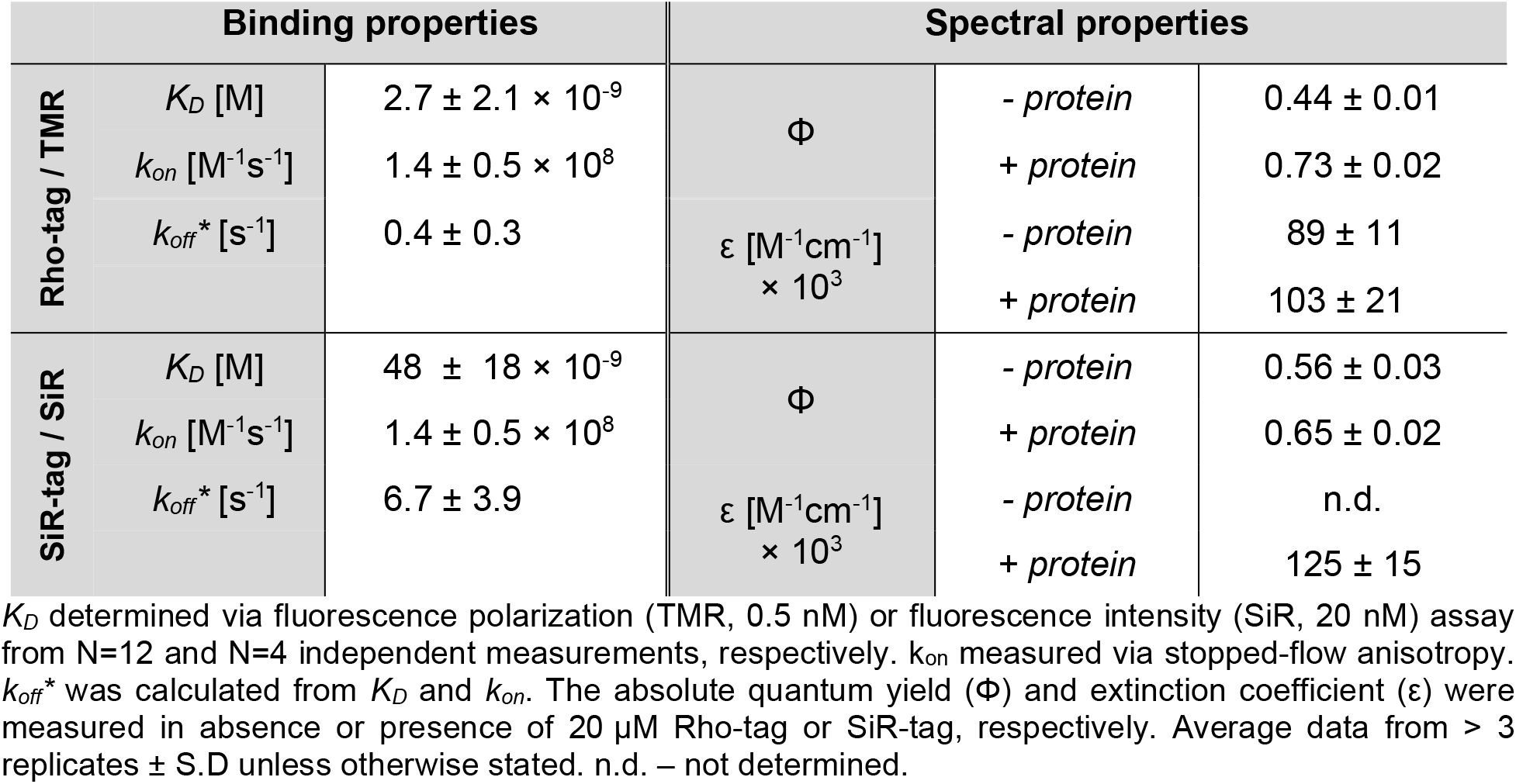
Binding and spectral characterization of the Rho-tag and SiR-tag system.

**Figure 2.**
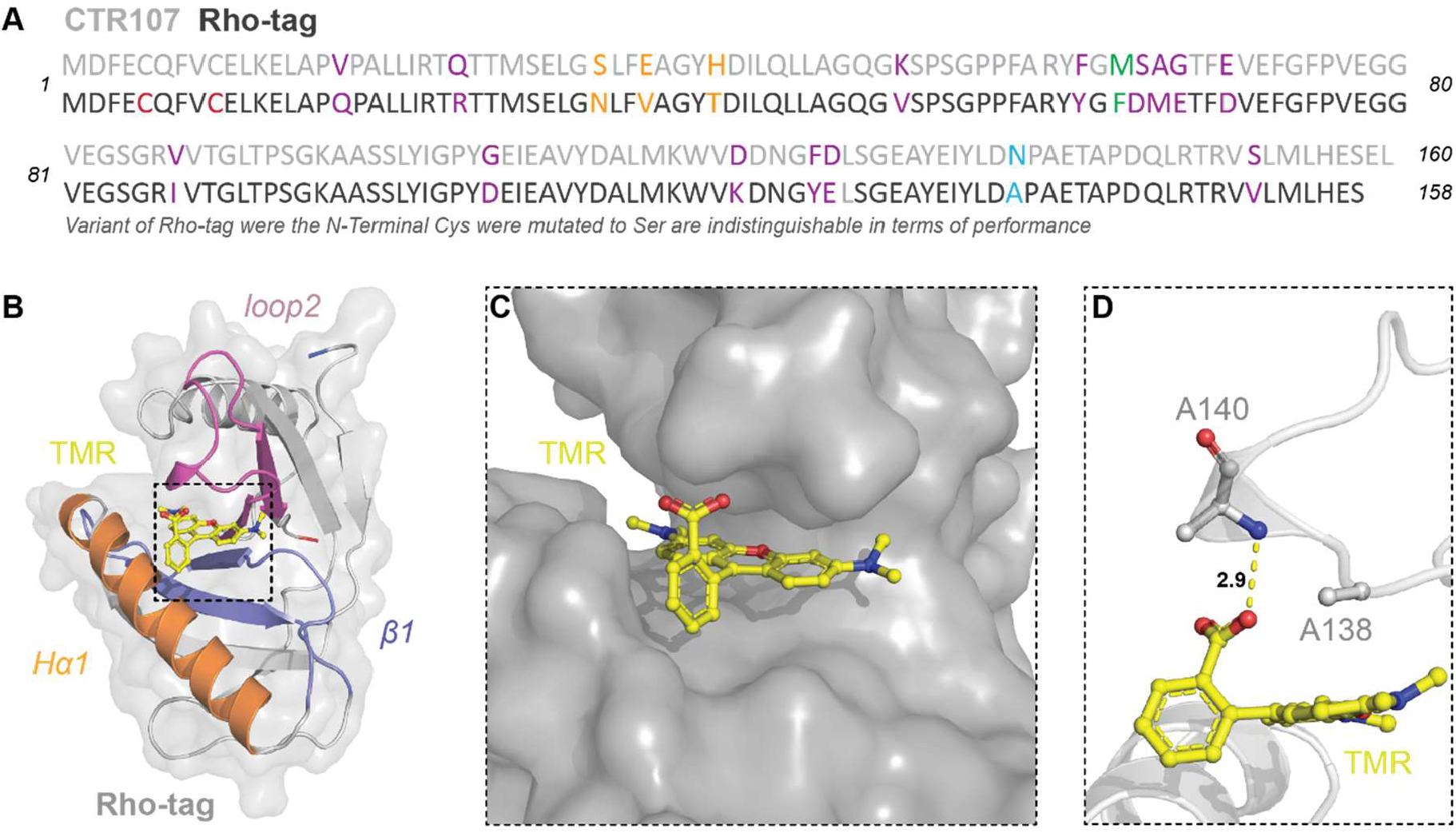
Structural analysis of Rho-tag. A) Amino acid sequence alignment of CTR107 (light gray) and Rho-tag (dark gray). Amino acid number indicated. The two proteins share 87% of their sequence. Origin of mutations marked by color: N138A from rational engineering – blue, M64F from deep mutational scanning – green, PROSS design 4 – purple, Helix α1 – orange, N-terminal cysteines – red. N-terminal cysteines (5, 9) can be replaced by serines without compromising Rho-tag function. B) Crystal structure of Rho-tag in complex with TMR (PDB ID 9RTM, 2.1 Å resolution). Tertiary structures are represented as colored cartoons and surface. Secondary element sites of engineering colored according to legend. TMR ligand presented as yellow sticks. C) Surface representation of Rho-tag bound to TMR ligand (sticks) shows insertion of xanthene ring in binding pocket. D) Hydrogen bonding between the TMR *ortho*-carboxylate and the α-amino group of A140. The space needed to bind TMR tightly was created by the N138A mutation. Distance in Å.

Rho-tag could be labeled with TMR in live U2OS cells fused to H2B. The high affinity for TMR allowed the use of dye concentrations as low as 50 nM. The signal-to-background ratio (S/B) was similar to that observed for HaloTag7 (Figure 3A). Live-cell labeling, as well as pre- and post-fixation staining, all yielded bright signals (Figure 3B). Washing gradually removed the bound rhodamine, demonstrating the reversibility of the labeling (Figure 3C). Full labeling of nuclear localized Rho-tag (NLS_3_) in live cells was reached within a few seconds (Figure 3D, E), whereas complete labeling of HaloTag7 under the same conditions required approximately 10 minutes. We attribute the faster live-cell labeling of Rho-tag, relative to HaloTag, to its superior labeling kinetics^[21]^ and the improved permeability of TMR compared to TMR-HTL. Non-covalent labeling strategies can help to reduce photobleaching by the exchange of damaged fluorophores.^[22]^ However, bleaching experiments with nuclear Rho-tag did not show a significant improvement in the apparent photostability compared to covalently labeled HaloTag7 (Figure S8).

**Figure 3.**
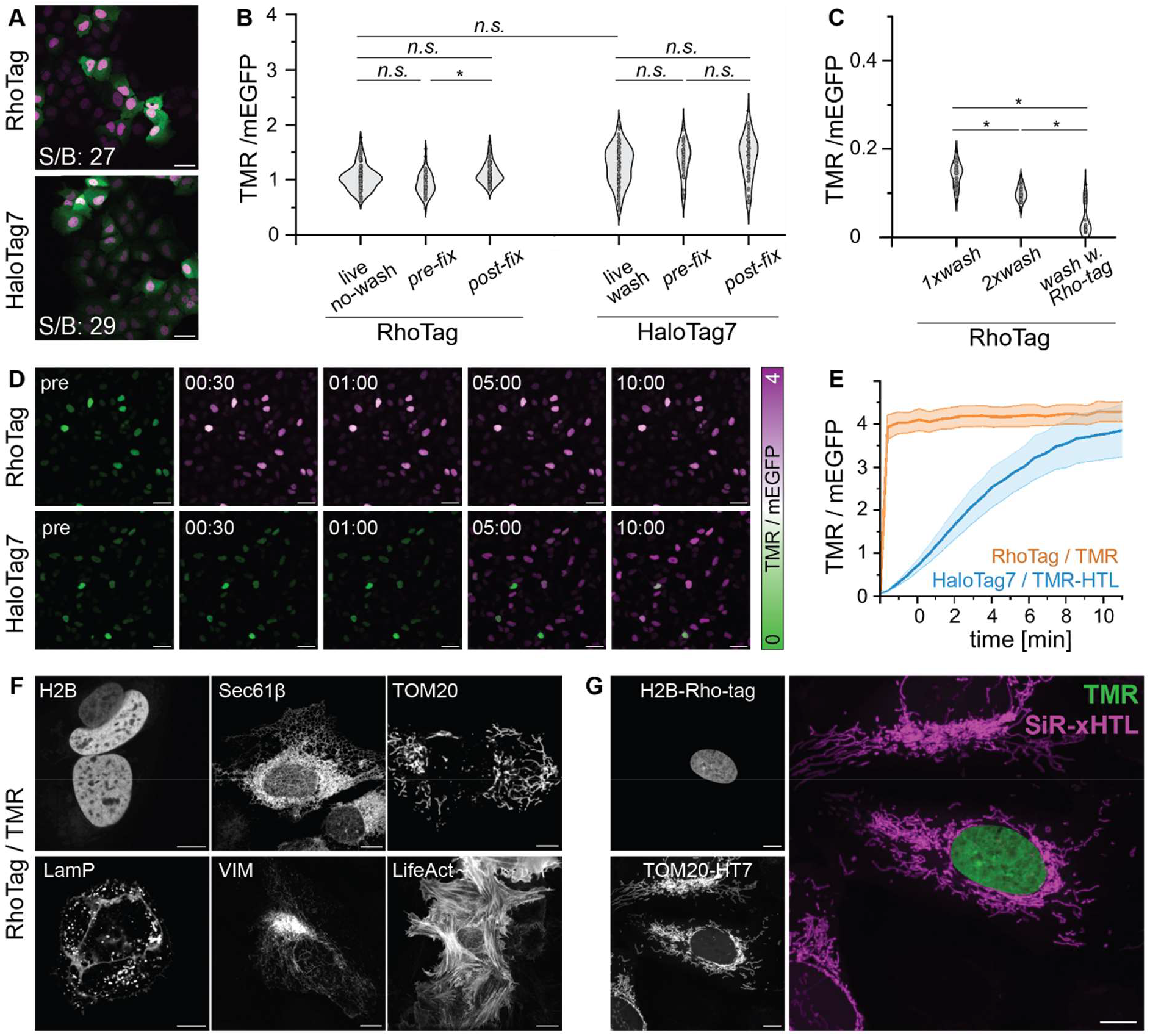
Live-cell labeling characterization of Rho-tag with TMR. A) Fluorescence micrographs (max. projections) of live U2OS cells expressing H2B-Rho-tag or -HaloTag7 labeled with TMR or TMR-HTL, respectively (50 nM). Co-translational mEGFP expression was used to identify transgene-expressing cells. Signal-over-background (S/B) ratios calculated as the average nuclear fluorescence intensity from cells expressing Rho-tag / HaloTag7 over non-expressing cells. Scale bars: 10 μm. B) Cellular intensity comparison between Rho-tag and HaloTag7 labeling with TMR in live-cells or under different fixation conditions extracted from similar images as in A). >100 nuclei were quantified from 3 images, global background subtracted and TMR signal divided by the individual mEGFP signal. Significance was calculated using two-sided t-tests including the Welch correction. n.s.: p ≥ 0.05, *: p < 0.05. C) Cellular intensity comparison of Rho-tag under different washing conditions. Quantification as in B). Washing with Rho-tag was done with 50 μM purified protein in medium to scavenge TMR. D) Live-cell labeling kinetics of Rho-tag and HaloTag7. U2OS cells expressing Rho-tag- or HaloTag7-P_30_-mEGFP-NLS_3_ labeled with TMR or TMR-HTL (50 nM), respectively. Scale bars: 50 μm. E) Signal and SEM from experiments as shown in C. Quantified from > 60 nuclei normalized to the mEGFP signal. F) Overexpression (CMV promotor) of Rho-tag fusions to the following marker proteins: endoplasmic reticulum – Sec61β, outer mitochondrial membrane – TOM20, lysosomes – LamP1, intermediate filaments – Vimentin (VIM), actin - LifeAct. Max. projections. Scale bars: 50 μm. G) Live-cell confocal multiplexing of H2B-Rho-tag / TMR (50 nM, nucleus, stable cell-line) and TOM20-HaloTag7 / SiR-xHTL (500 nM, mitochondria, rAAV). Max projection. Scale bar: 10 μm.

Rho-tag could be localized to various cellular structures (histones, endoplasmic reticulum, mitochondria, lysosomes, intermediate filaments, and actin) in live cells (Figure 3F). Finally, we investigated if Rho-tag can be multiplexed with established labeling strategies for derivatized rhodamines. Linking an exchangeable HaloTag Ligand (xHTL)^[22]^ *via* carbon 6 of the benzyl ring of TMR reduced the binding affinity to Rho-tag by more than 7000-fold (*K*_*D*_ 18.8 ± 7.4 µM) which enabled imaging of both tags with rhodamines within one cell (Figure 3G).

### Circular-permuted and bioluminescent Rho-tag variants

The original *N*- and *C*-termini of Rho-tag are relatively distant from the fluorophore binding site (26 and 30 Å, respectively). Having the termini close to the site of the fluorophore can be advantageous for the generation of biosensors.^[23]^ To achieve this, we connected the original termini with a designed linker, using RFdiffusion for backbone generation and ProteinMPNN for sequence design^[24-26]^, and introduced new termini at positions 64/67 (supplementary note 4). This yielded cpRho-tag which offered high thermostability (60 °C) and retained its high binding affinity to TMR (*K*_*D*_ 6.8 ± 5.8 nM). In live-cell labeling experiments, cpRho-tag performed as well as Rho-tag (Figure S9).

Bioluminescence reporters based on bioluminescence resonance-energy transfer (BRET) have gained popularity in bioimaging and -biosensing due to their high sensitivity.^[27]^ We either fused NanoLuc (NLuc) to the *N*- or *C*-terminus of Rho-tag or inserted a circular-permuted version of NLuc^[28]^ at position 64/67 of Rho-tag to create RhoLuc (Figure 4A). The chimeric RhoLuc displayed four-times higher BRET efficiency than the terminal fusions (Figure 4B).

**Figure 4.**
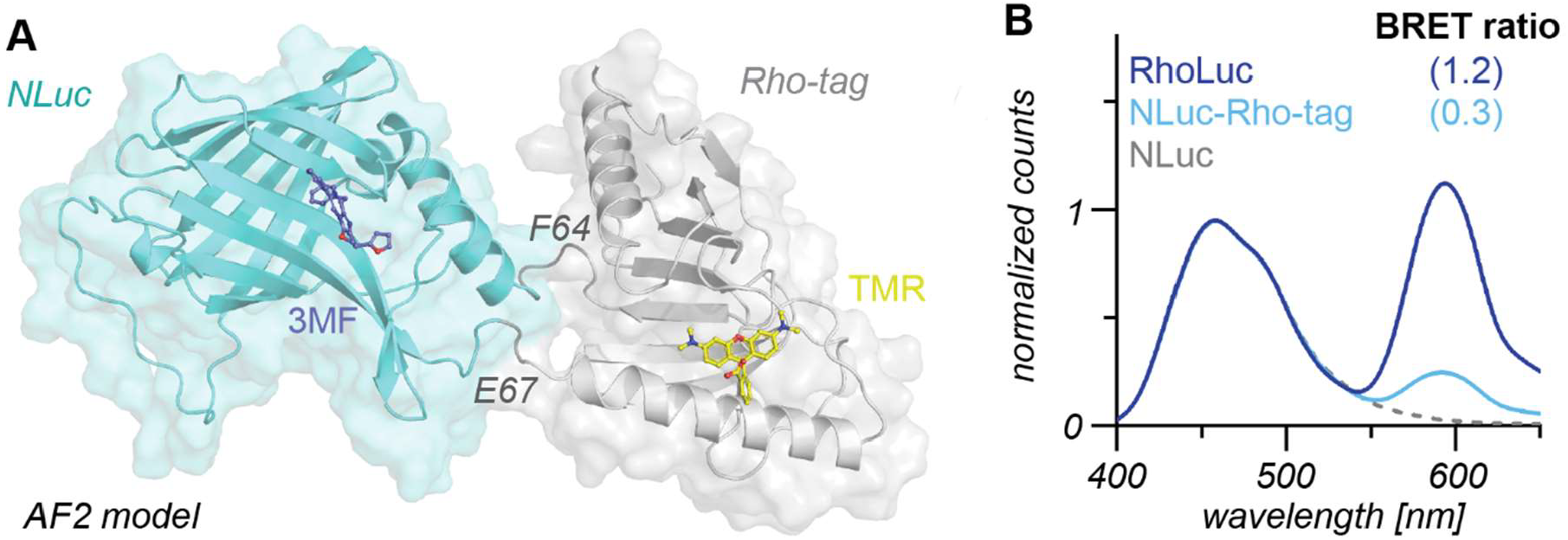
Chimeric RhoLuc enables BRET. A) AlphaFold2 (AF2) model of RhoLuc (*N*-Rho-tag^1-64^-GSG-cpNLuc^67/68^-GS-Rho-tag^67-158^-*C*). cpNLuc^67/68^ was inserted into 64/67 site of Rho-tag. Tertiary structures are represented as colored cartoons and surface. TMR ligand presented as yellow sticks. 3-methoxyfurimazine (3MF) was modeled into the structure by aligning a crystal structure of NLuc (PDB ID 7SNT) to the AF model and is represented as blue sticks. B) Bioluminescence profil of RhoLuc (dark blue, insertion), NLuc-Rho-tag (light blue, terminal fusion) or NLuc (gray). Proteins (2 nM) were labeled with excess TMR (50 nM). The fluorescence emission spectra were recorded upon addition of Nano-Glo^®^ substrate (Promega 1:1000). BRET ratio given as I_593_/I_459_ nm. Average data from three technical replicates.

### SiR-tag - a fluorogenic binder for far-red emitting dyes

Silicon rhodamine^[29]^ is a commonly used fluorophore for bioimaging, due to its far-red emission wavelength and excellent fluorogenicity. However, Rho-tag displayed only weak binding affinity for SiR derivatives (*K*_*D*_ 6.5 ± 2.1 µM). We therefore used Rho-tag as a platform to develop a silicon rhodamine binding protein tag (SiR-tag) *via* a two-step directed evolution strategy.

First, we randomized positions 133 to 149 close to the xanthene (*loop2*, Figure 2B) by creating libraries with four randomized codons each (supplementary note 5). We submitted the combined libraries to yeast display and five rounds of FACS screening which yielded eight putative SiR-tag candidates. Amongst them, the variant with mutations A138S / P139R / A140H showed a 72-fold improved binding affinity for the SiR derivative JF_646_^[20]^ (*K*_*D*_ 92 ± 35 nM). We increased the thermostability of the protein using PROSS and further randomized positions 51 to 79 (*β1*, Figure 2B), following a similar strategy as described above. This yielded SiR-tag (Figure 5A) which displayed high binding affinity to JF_646_ (*K*_*D*_ 20 ± 14 nM) and SiR (*K*_*D*_ 48 ± 18 nM). SiR-tag displays high stability (*T*_*M*_ 72 °C) and fast exchange kinetics, similar to Rho-tag (Table 1). Moreover, SiR exhibited a strong fluorescence increase (13-fold) upon SiR-tag binding, which is comparable to the covalent labeling of HaloTag7 with the same fluorophore (Figure 5B). SiR-tag has an 87-fold higher binding affinity for JF_646_ over TMR, whereas Rho-tag has 2,400-fold higher binding affinity for TMR compared to JF_646_. The high binding specificities of Rho-tag and SiR-tag for their respective substrates might allow use of this protein pair for multiplexing.

**Figure 5.**
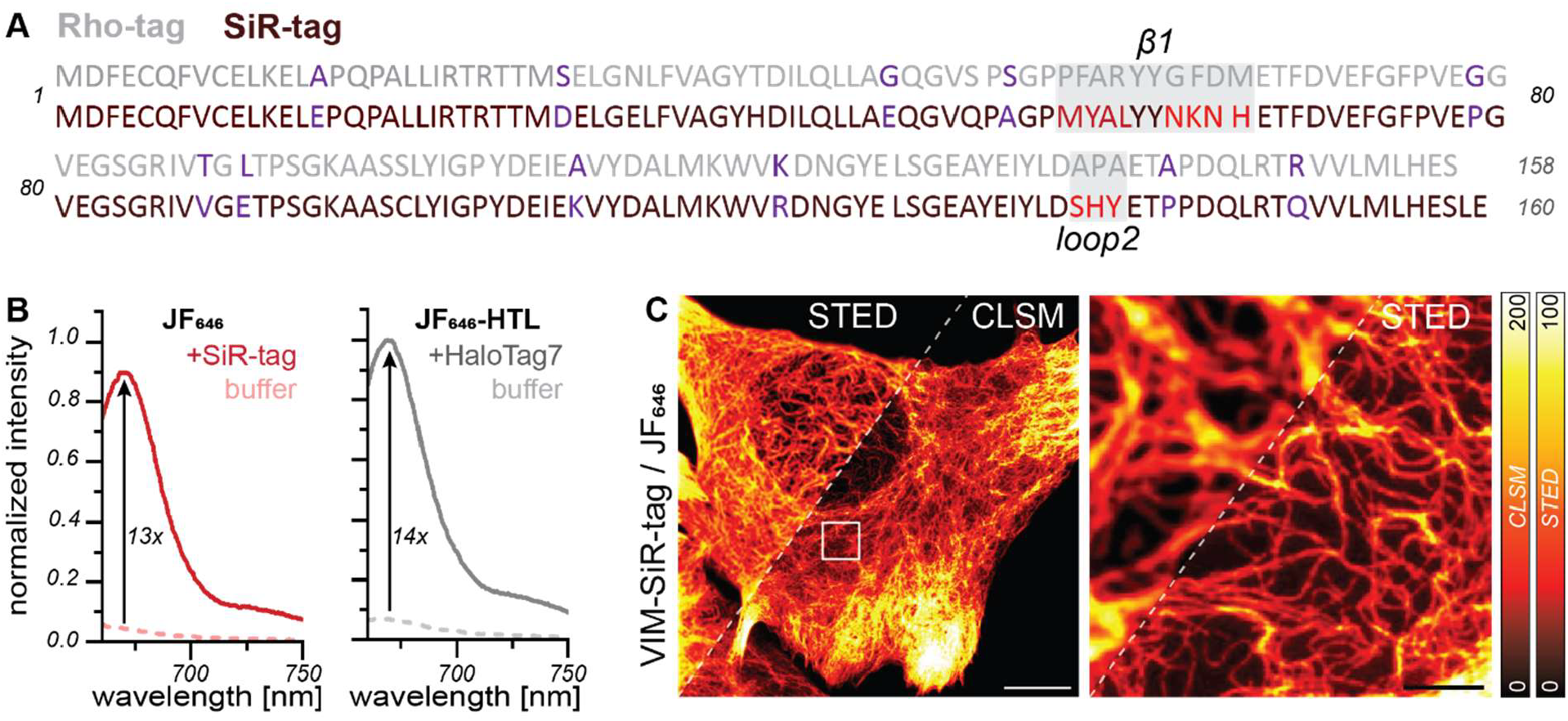
SiR-tag – a fluorogenic binder for far-red emitting dyes. A) Amino acid sequence alignment of Rho-tag (light gray) and SiR-tag (dark red). Amino acid number indicated. The two proteins share 82% sequence identity. Origin of mutations marked by color: PROSS design 8 – purple, Helix *α1* – orange, loop2 and β1 directed evolution – red. B) Fluorescence emission increase of SiR upon binding to SiR-tag and covalent binding to HaloTag7. The fluorescence emission spectrum of 50 nM dye was recorded in presence and absence of 20 µM protein. F_I_/F_I0_ indicated by black arrow. Average data from technical triplicates. C) Live-cell CLSM and STED microscopy of U2OS cells expressing VIM-SiR-tag and labeled with JF_646_ (500 nM). Scale bars: 10 µm (overview) and 2 µm (magnification). Pixel intensities scaled according to reference bar.

SiR-tag could be labeled with JF_646_ in live U2OS cells expressing H2B-SiR-tag. The staining revealed clear nuclear localization at low dye concentrations (200 nM) with outstanding signal-to-background, even though the labeling was 1.5-fold less bright compared to covalent HaloTag7 labeling (Figure S10). Furthermore SiR-tag was compatible with live-cell STED microscopy of intermediate filaments (Figure 5C).

### Rho-tag and SiR-tag for advanced microscopy

The reversible yet fast binding and unbinding of the Rho-tag system makes it attractive for SMLM (Figure 6A).^[12]^ First, we determined the binding time of both Rho-tag and SiR-tag in single-molecule imaging experiments conducted in U2OS cells expressing the monomeric membrane receptor CD86 fused to Rho-tag or SiR-tag. At low dye concentrations (3 nM), recurring single-molecule binding events to single CD86-Rho-tag proteins were detected (Video S1), which allowed us to resolve single CD86 clusters (Figure 6B) and extract binding times of 1019 ± 76 ms (Rho-tag) and 345 ± 45 ms (SiR-tag), respectively. To demonstrate the use of Rho-tag as a tool for super-resolution imaging of subcellular structures, we resolved the intermediate filaments with a resolution of 21 nm in chemically fixed U2OS cells expressing Vimentin-Rho-tag (Figure 6C). Similarly, we performed MINFLUX imaging to resolve single Vimentin molecules along the intermediate filaments in fixed cells (Figure 6D).

**Figure 6.**
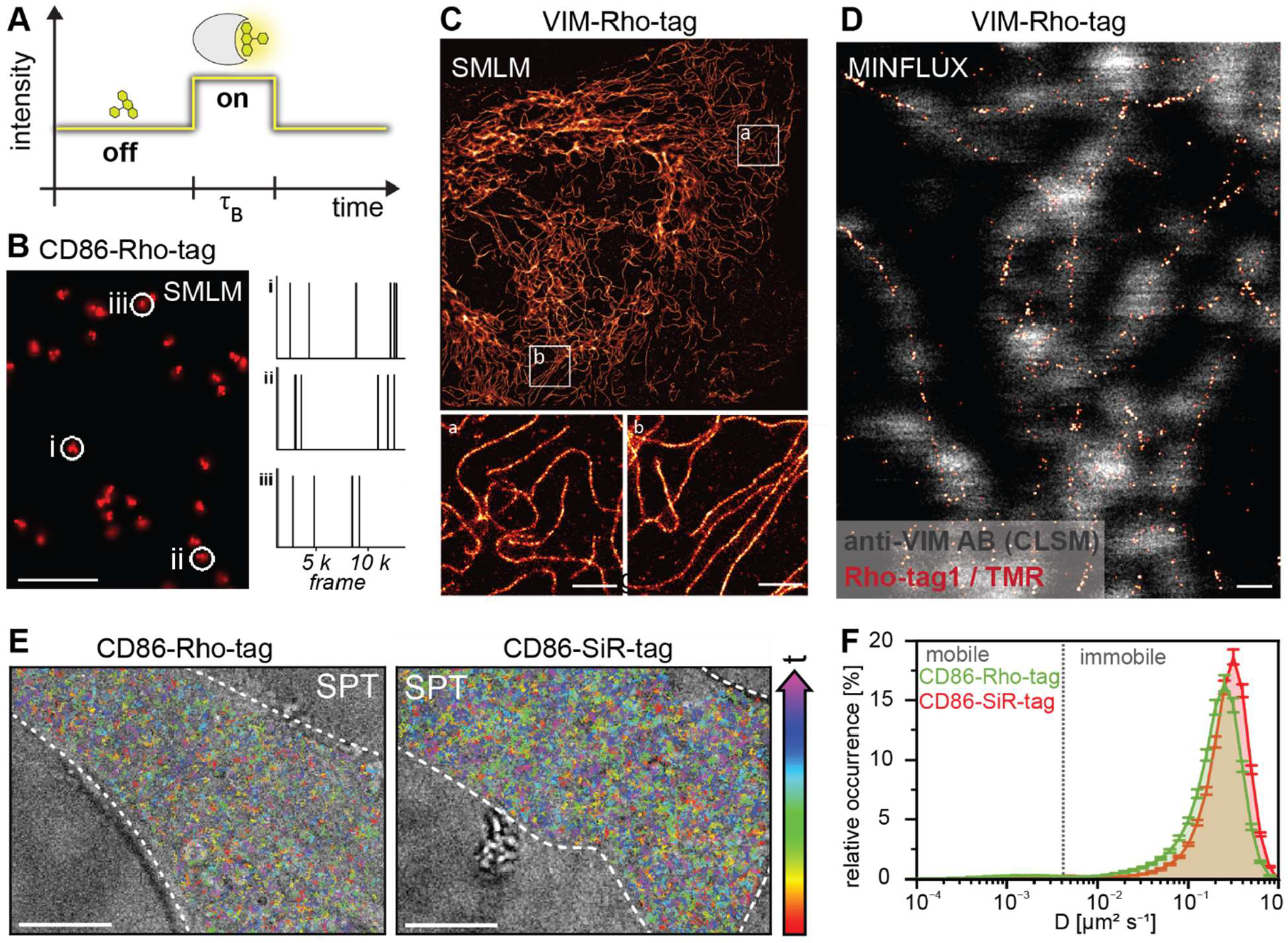
Rho-tag applications for single-molecule localization microscopy. A) Rho-tag-SMLM imaging concept. Transient binding of TMR to Rho-tag induces local signal spikes (single-molecule traces) during the bright time τ_B_ and until spontaneous unbinding of the dye. B) SMLM imaging with TMR (3 nM) on fixed U2OS cells resolved single CD86-Rho-tag clusters. Scale bar: 0.5 µm. Single-molecule traces of highlighted clusters yield a bright time of 1,019 ±76 ms. C) Super-resolution reconstruction of Vimentin filaments (25,000 frames, 150 ms exposure time) using Rho-tag and TMR (1 nM) in U2OS cells. The resolution was determined using decorrelation analysis^[41]^ (21 nm). Scale bars: 10 µm (overview) and 2 µm (magnification). D) 2D-MINFLUX imaging (100 photons last iteration, 2 h) with TMR (0.2 nM) in U2OS Vim-Rho-tag cells with a localization precision of 2.2 nm. Co-staining of Vimentin with primary anti-Vimentin and secondary AF647-labeled antibodies. Scale bar: 0.2 µm. E) Single-particle tracking (SPT) in living U2OS cells. Single-molecule trajectories recorded of TMR binding to CD86-Rho-tag (left) or SiR binding to CD86-SiR-tag (right) expressed in U2OS cells. Trajectories are color-coded by their time of occurrence as indicated by the colored arrow: early appearing trajectories denoted in red and late appearing trajectories colored in purple. Mean trajectory lifetime per cell (0.65 ± 0.01 s) at a laser power of 0.5 kW/cm^2^. Scale bar: 10 µm. F) Histogram of relative abundance of diffusion coefficients reveals immobile (4.3 ± 0.2%) and mobile (95.8 ± 0.2%) trajectories among all cells (N = 152). Error bars represent SEM.

Live-cell super-resolution imaging with high temporal resolution can be achieved by combining exchangeable probes and high-density single-molecule localization using a neural network^[30]^ which was shown to resolve cellular features under physiologically relevant time-scales.^[31]^ We trained a DeepSTORM^[32]^ model with single-molecule data of Rho-tag-Sec61β and used this model to reconstruct the movement of the ER by live-cell SMLM (Video S2, Figure S11).^[31]^ Another powerful single-molecule imaging method in living cells is single-particle tracking. The integration of exchangeable probes has recently been shown to report diffusion coefficients and dynamic clustering of membrane proteins in long-term experiments without substantial photobleaching.^[33]^ Using living U2OS cells expressing CD86 fused to Rho-tag or SiR-tag (Figure 6E), a high number of single-molecule trajectories was recorded in comparison to non-transfected U2OS cells (Figure S12A), of which > 95% were classified as mobile by their diffusion coefficient (Figure 6F). The average trajectory lifetime was ∼0.6 s for both protein tags, which is comparable to values reported for non-covalent labels (Figure S12B).^[33]^ Over the imaging course of 30 min, we observed a signal reduction of ∼30% for Rho-tag and ∼50% for SiR-tag (Figure S12C), demonstrating the suitability of both tags for long-term single-particle tracking in living cells.

### In vivo brain imaging of zebrafish larvae

TMR and SiR are predicted to readily cross the blood-brain barrier (Figure 1A). This prediction is supported by their instant live-cell labeling kinetics (Figure 3D). We investigated the labeling of Rho-tag and SiR-tag expressed in neuronal cells of zebrafish larvae, a popular model organism in biology.^[34]^ The staining of HaloTag7 in zebrafish larvae requires long incubation times (hours) and relatively high dye concentrations (4 – 30 µM).^[35-37]^

Labeling of Rho-tag in neurons of live zebrafish larvae (3-4 *dpf*, mosaic expression) was observed within 1 hour using dye concentrations as low as 250 nM (Figure 7A, B). In contrast, HaloTag7 could not be labeled under these conditions using the same dye as an HTL derivative (Figure 7A, B), and required overnight staining at 10 µM dye concentration for effective staining (Figure S13). Imaging of zebrafish larvae expressing Rho-tag and labeled with the TMR derivative JF_549_ was done in the presence of the dye. However, most signal persisted when the larvae were washed and left freely swimming for 10 minutes in water not containing JF_549_ (Figure 7D). Similarly, SiR-tag enabled efficient labeling in neurons of live zebrafish larvae. In contrast, HaloTag7 labeling was less efficient under equivalent conditions (Figure 7E). These experiments demonstrate the potential of Rho-tag and SiR-tag for *in vivo* imaging.

**Figure 7.**
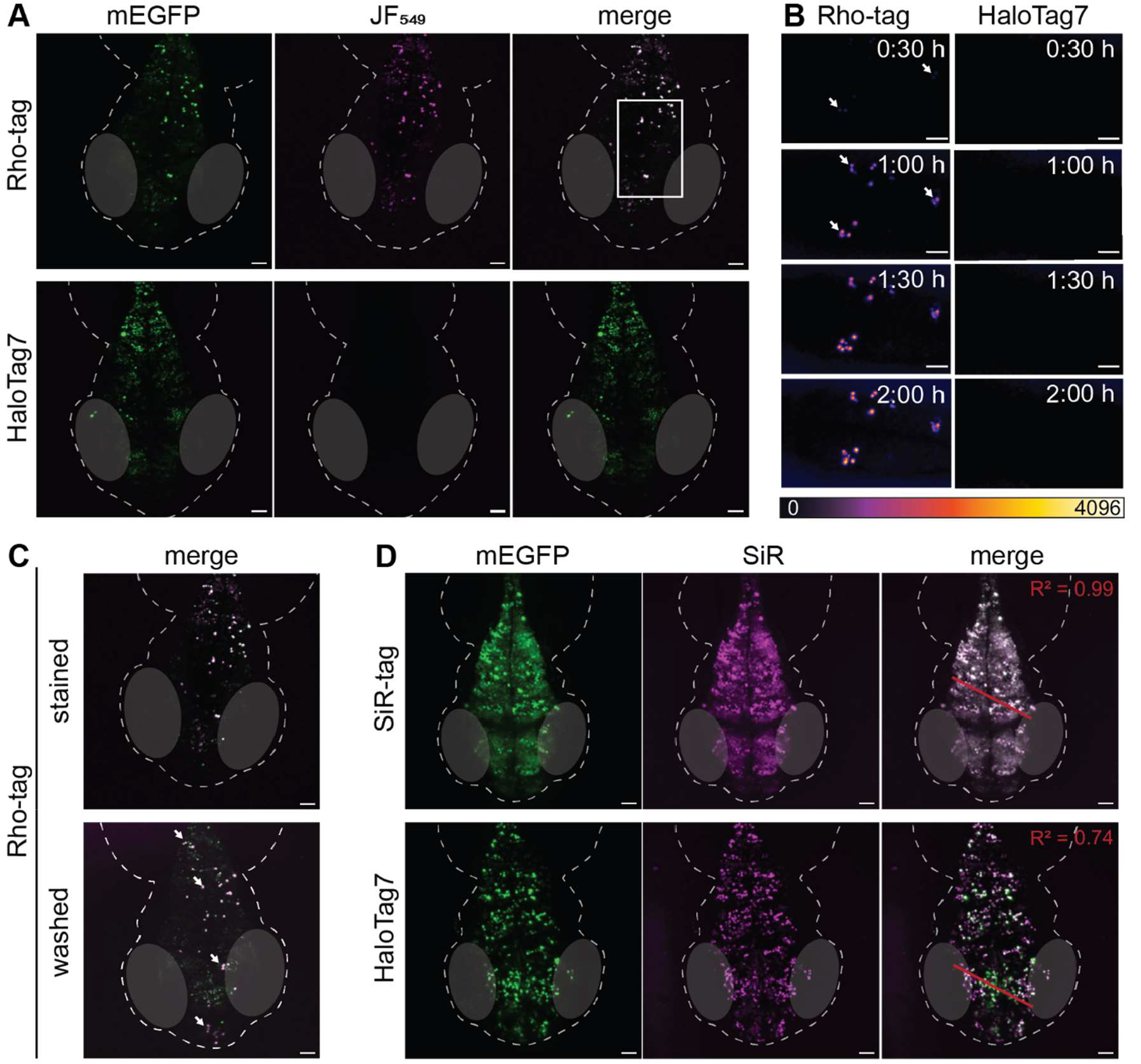
Neuronal labeling of Rho-tag and SiR-tag in live zebrafish larvae. A) Fluorescence micrographs (max. projections) of zebrafish embryos mosaically expressing neuronal Rho-tag- (top) or HaloTag7-P_30_-mEGFP-NLS_3_ (bottom) with the TMR derivative JF_549_ or its respective HaloTag Ligand (250 nM, 2 h). Live larvae were embedded in agarose (supplemented with Tricaine) and imaged on a confocal microscope over 200 μm depth. Scale bars: 50 μm. N=6 (Rho-tag) and N=8 (HaloTag7) individual larvae delivered similar results. B) Live *in vivo* labeling kinetic comparison of Rho-tag and HaloTag7 probes. Inset from (A). Larvae were embedded in agarose and 250 nM dye added to the medium. White arrows show onset of labeling signal. Scale bars: 10 μm. C) Comparison of Rho-tag labeling under no-wash and wash conditions in live zebrafish embryos. Larvae expressing neuronal Rho-tag were labeled in agarose as described in (A) and (B) and afterwards freed, washed twice, left swimming in fresh water for 10 min and imaged again. White arrows exemplarily indicate remaining signal after wasg. Merge of JF_549_ – magenta, mEGFP – green. Scale bars: 50 μm. D) Comparison of SiR-tag and HaloTag7 labeling in live zebrafish embryos. Live confocal imaging of zebrafish embryos expressing neuronal GOI-P_30_-mEGFP-NLS_3_ (mosaic) labeled with SiR or its respective HaloTag Ligand (250 nM, 2 h). Max. z-projection (200 μm). Scale bars: 50 μm. Efficiency of labeling was investigated by the colocalization between SiR and mEGFP signals along the red line and characterized by the R^2^. N=3 (both SiR-tag and HaloTag7) individual larvae showed similar results.

## Discussion

We introduce two new fluorophore-binding proteins, Rho-tag and SiR-tag, which bind to unsubstituted (silicon-) rhodamines with high binding affinities (nanomolar *K*_*D*_). Both tags were derived from a bacterial multidrug-resistant protein with a weak intrinsic affinity for rhodamines. Through protein engineering, the affinity of their predecessor towards the fluorophores were increased by about three orders of magnitude, with a *K*_*D*_ of Rho-tag for TMR of about 3 nM. The rate of substrate binding to the two tags is close to be diffusion controlled. The crystal structure of Rho-tag bound to TMR reveals that the xanthene core is bound deep in the binding pocket of the protein, helping to rationalize the two-fold increase in brightness of TMR when bound to Rho-tag. The structural information on the dye-protein interaction will aid in future engineering of the tags. Rho-tag and SiR-tag are small and stable single-domain proteins with about 18 kDa size and a melting point of 61 and 72 °C, respectively. Rho-tag can be subjected to circular permutation and allows insertion of other proteins into its structure. For example, we have inserted the luciferase NanoLuc into a loop of Rho-tag such that the fluorophore is close to the active site of the luciferase, creating a chimera with relatively high BRET efficiency. These experiments suggest that Rho-tag, SiR-tag and their circular permutated variants represent an attractive platform for the design of biosensors.

The main applications of Rho-tag and SiR-tag are in live-cell imaging. We demonstrate that Rho-tag expressed in mammalian cells can be labeled with TMR within seconds. Labeling of Rho-tag is significantly faster than the labeling of self-labeling protein tags such as HaloTag, which we attribute at least partially to the higher cell permeability of unsubstituted TMR relative to that of the TMR derivate used for HaloTag labeling. Rho-tag can also be labeled in fixed samples. Importantly, Rho-tag and SiR-tag are compatible with various super-resolution microscopy techniques, including STED, SMLM, and MINFLUX, and shows excellent performance in single-particle tracking. Furthermore, as Rho-tag and SiR-tag show very low affinity towards the substrates of the self-labeling tags HaloTag and, presumably, SNAP-tag, they can be used in combination with these tags.

The labeling of Rho-tag is reversible and the fluorophore can be washed out after labeling. For most applications, imaging of Rho-tag and SiR-tag fusion proteins is thus done in the presence of the added dye. The high affinity of the tags for their substrates, which allows their use at nanomolar concentrations, and the fluorogenicity of the used dyes, in particular of SiR and its derivatives, enable live-cell confocal imaging without significant background. As binding of the dyes to Rho-tag and SiR-tag affects their fluorescence lifetimes, a further reduction of background signal from unbound dye could potentially be achieved through fluorescence lifetime imaging microscopy.^[38]^ We note that the reversible binding of TMR to Rho-tag did not result in an increase in photostability under strong laser intensities when compared to HaloTag7-bound TMR. We speculate that the tight binding affinity might prevent efficient exchange of damaged fluorophore or that the close proximity of the dye to the protein might result in photodamage to the protein.

The main motivation for the development of Rho-tag and SiR-tag was the assumption that the high permeability of unsubstituted rhodamines should facilitate labeling in complex samples, such as the central nervous system of a living animal. Indeed, Rho-tag expressed in neurons of zebrafish larvae could be efficiently labeled at substrate concentrations at which no labeling of HaloTag7 was observed. The signal of JF_549_-labeled Rho-tag persisted even after placing the labeled larvae in water in absence of the dye. Apparently, the wash-out of these dyes is slow enough to enable persistent staining in zebrafish larvae. Collectively, these experiments highlight the potential of Rho-tag and SiR-tag for *in vivo* applications.

Rho-tag and SiR-tag are part of a new class of tags that reversibly bind to unsubstituted rhodamines. The other members of this class are *de novo* designed rhodamine binders (Rhobins)^[39]^, which have similar features such as Rho-tag and SiR-tag, and which have been developed in parallel to this work. These rhodamine-binding tags complement tags such as FAPs and the FAST system, which are engineered proteins that bind to different types of fluorogens. Another system for reversible fluorescence labeling of proteins is the labeling of HaloTag7 with xHTLs^[22]^. The distinguishing feature of the rhodamine-binding tags relative to these other tags for applications in live-cell imaging is their outstanding spectroscopic properties such as brightness combined with the excellent permeability of their substrates.

## Supporting information

Supplementary informations

## Acknowledgement

Diffraction data were collected at the Swiss Light Source, beamline X10SA, of the Paul Scherrer Institute, Villigen, Switzerland. The authors are grateful to J. Wittbrodt, J. Benjaminsen and T. Thumberger (COS Heidelberg) for providing resources, infrastructure and support for experiments with zebrafish larvae. We thank J. Reinstein (MPIMR) for giving us access to a stopped-flow kinetics device. The authors thank B. Koch, A. Bergner, J. Kress, P. Breuer, A. Herold, V. NasufoviĆ, B. Réssy, and D. Schmidt (all MPIMR) for providing reagents or material. We thank the mass spectrometry facility of the MPIMR (S. Fabritz, T. Rudi, and J. Kling) and C. Room (IT MPIMR) for their support, M. Siegfried (Goethe University Frankfurt) for his assistance in implementing RhoTag-SMLM, and M. Scherer (Fleishman lab, Technion) for his assistance with PROSS. The authors were supported by the Heidelberg Biosciences International Graduate School (HBIGS), the International Max-Planck Research School on Cellular Biophysics (IMPRS-CBP) and the Max Planck School Matter to Life supported by the German Federal Ministry of Education and Research (BMBF) and in collaboration with the Max Planck Society.

