## Supplementary informations for "Fast, Bright and Reversible Rhodamine Tags for Live-Cell Imaging"

### Supplementary notes

#### Note 1: directed protein engineering of Rho-tag

The directed protein evolution workflow is shown in Figure S6A. For Rho-tag, a synthetic deep mutational scanning library (sDMSL) was based on CTR107$^{N138A}$. Within this library, all 160 amino acids were individually replaced with all other 19 amino acids, not considering cysteines, or deletions, yielding a theoretical size of 3.2 × $10^3$. Based on the crystal structure of CTR107$^{N138A}$ (PDB ID 9RTL) we reasoned, that *Helix α1 (Hα1)* interacts with the lower benzyl ring of TMR and the *ortho*-carboxy moiety (Figure S6B). Site-saturation mutagenesis libraries (SSMLs) containing three randomized codons at a time (Supplementary Table 4) were prepared for *Hα1*, yielding a library diversity of 9.3 × $10^3$ per library. The single SSMLs were pooled (total diversity 1.9 x$10^5$). All DNA libraries were designed for subsequent homologous recombination (HR) with a linearized donor vector into yeast and transformed with a transformation efficiency covering the theoretical library size by at least two orders of magnitude.

For subsequent screening, the yeast displaying the randomized libraries were labeled with MaP555[1], a fluorogenic TMR analogue with reduced background signal. Selecting a high-affinity MaP555 binder was not possible, however, the strong selection pressure helped to co-evolve high-affinity binder for TMR. Throughout FACS sorting, the dye was present within the medium at decreasing fluorophore concentrations (500 – 10 nM). We collected the top 1 – 10% brightest yeast cells to enrich variants with putative better binding or spectral properties. The selection process was monitored by a shift of the whole yeast population towards brighter labeling. Next-generation sequencing results were analyzed by calculating the sequence frequencies (number of each sequence over total amount of analyzed reads) which revealed a high diversity of the unsorted library, with a minor parental bias (2.3%). H40Y / Q44P / L45P / L46Q enriched strongly but was not stable enough to be recombinantly expressed. The M64F mutation was strongly enriched from the sDMSL (from 5% to 79%) but only had little impact on the binding affinity and the protein stability.

#### Note 2: improving the thermostability using PROSS

Rational engineering of Rho-tag0.1 caused a drop in melting temperature by -8.7° C ($T_M$ 59.6 to 50.9 °C). Most variants selected for better TMR binding revealed further decreased thermostabilities, potentially restricting the evolvability towards high-affinity rhodamine binding. Therefore, we fed the crystal structure of CTR107$^{N138A}$ (PDB ID 9RTL) into the Protein Repair One-Stop Shop[2]. PROSS found 251 phylogenetically related sequences, most of them having a good coverage (>90%) but little sequence identity (~40%). The PROSS designs are summarized in Supplementary Table 2. Recombinant expression and measurement of the thermostability of a selection of PROSS designs revealed a gradual improvement in thermostability up to 78° C (Supplementary Table 2), whereas 14 mutations (design 4) provided the best compromise between stability and function *i.e.* TMR binding and were therefore included into the Rho-tag final design. More challenging engineering efforts, like the development of SiR-tag and RhoLuc, required using 20 more mutations suggested by PROSS (design 8) in order to maintain high stability.

**Note 3: site-directed mutagenesis of *N*-terminal Cys residues**

The *N*-terminal disulfide bridge of Rho-tag (C5, C9) may restrict its application in biochemical assays containing thiol-reactive reagents or in cellular compartments having oxidative microenvironments. We therefore used site-directed mutagenesis to replace both cysteine residues with serine. Rho-tag$^{C5S,\ C9S}$ shows a decreased melting temperature (51 °C), however, the TMR binding affinity ($K_D$ 4.4 ± 1.6 nM) was indistinguishable from Rho-tag.

**Note 4: directed protein engineering of cpRho-tag**

For cpRho-tag engineering, we introduced the mutations suggested by PROSS design 8 (Supplementary Table 2) into Rho-tag to increase the protein stability and evolvability. To bridge the *N*-to-*C*-distance, we initially connected the original termini with a flexible $(GGS)_5$ linker and screened for new termini in solvent-exposed loop regions at the following positions: 50/52, 53/55, 54/56 64/67, 83/85, 140/142, 140/147, 141/143, 142/147, 142/146, 145/147. Among these, only the cp64/67 variant retained strong binding affinity for TMR and high thermostability. Moreover, we found that truncating the first 18 *N*-terminal residues and the final 3 *C*-terminal residues had no negative impact on Rho-tag function.

We next used computational tools to design an optimized linker to replace the flexible GGS linker, with the goal to generate sequences that participate in folding and feature interactions with the rest of the protein. This strategy has been successfully applied to other circularly permuted proteins to enhance stability and activity.[3] We generated 240 backbone models connecting the termini of the truncated protein (A19-M154) with linker lengths ranging from 8 to 15 residues using RFdiffusion[4] (default settings). For each linker backbone, a sequence was designed using ProteinMPNN[5], keeping the remainder of the protein sequence fixed and excluding cysteine from the set of designable residues. The structures of the resulting sequences were then predicted using ColabFold[6] (with five recycles and one amber relax step). Predicted models were ranked by the pLDDT confidence score of the linker region (Supplementary Table 3). Among the top candidates, six linker variants were experimentally tested (melting temperature, TMR binding affinity), with one sequence (LKNAGPETETLESQP) clearly superior to the others.

**Note 5: directed protein engineering of SiR-tag**

For the engineering of *loop2,* SSM libraries containing four randomized codons at a time (Supplementary Table 5) were prepared as described above having a theoretical library diversity of $2.0 \times 10^5$. Libraries were pooled (diversity $4.9 \times 10^6$) and enriched for 5 consecutive rounds of FACS using $JF_{646}$ (500 – 100 nM) for fluorescence labeling. From 14 analyzed single colonies, eight putative SiR-tag variants were found and tested *in vitro* (Supplementary Table 7). The variant containing A138S / P139R / A140H was combined with the PROSS design 8 (Supplementary Table 2). This new variant was used again as the template for the generation of SSM libraries on *β1* similar to *loop2* (Supplementary Table 6). From high throughput screening, six variants were tested *in vitro (*Supplementary Table 7*)*, of which P57M / F58Y / R60L and G63N / F64K / D65N / M66H were combined yielding SiR-tag.

### METHODS

#### 1. Computational methods

*Physicochemical analysis*

Pharmacokinetic analysis was performed using the SwissADME webtool.[7, 8] Compounds structures were used as SMILES as input to generate BOILED-Egg[9] plots and physicochemical properties.

*PROSS*

The webpage-based "Protein Repair One-Stop Shop" (PROSS) was used to generate Rho-tag designs offering increased thermostability. The crystal structure of CTR$^{N138A}$ (PDB ID 9RTL) was used as input and without any other constrains.[2, 10]

*Alphafold*

Protein structure predictions were generated using AF2 in multiple sequence alignment mode from the primary sequence[11] on the *Raven* supercomputer (MPG).

*Molecular dynamics simulation*

Molecular dynamics (MD) simulations were carried out for 200 ns in Maestro (Schrödinger Inc., New York) under constant pressure (NPT ensemble) to evaluate the interactions between TMR ligands and Rho-tag or HT7. The protein–ligand complexes were obtained by X-ray crystallography (Rho-tag – PDB ID 9RTM; HaloTag7 PDB ID 7ZIY), and prepared using the Protein Preparation Wizard with default settings. An orthorhombic periodic box with a 10 Å buffer distance was constructed using the Desmond System Builder module. Sodium and chloride ions were added to neutralize the system and replicate an experimental salt concentration of 0.15 M NaCl. The simulation system, was parameterized using the OPLS4 force field. MD simulations were conducted using the default algorithms and protocols of the Desmond Molecular Dynamics module. The system was equilibrated before executing a 200 ns production run at 300 K and 1.01325 bar.

#### 2. Chemical Procedures

TMR, $JF_{549}$, MaP555, SiR and $JF_{646}$ were synthesized following standard protocols.[1, 12-14] Custom-synthesized fluorophores were validated using HRMS (Method Table 1) on a Bruker maXis IITM ETD spectrometer with electron spray ionization (ESI) by the Mass Spectrometry facility (MPIMF, Heidelberg). 6-carboxy rhodamine HaloTag Ligands of TMR, SiR, $JF_{549}$ and $JF_{646}$ were synthesized by D. Schmidt and B. Réssy (MPIMF, Heidelberg) according to literature procedures.[12, 15, 16] TMR- and SiR-xHTL (S5) were used from to prior studies.[17] All other dyes were purchased from commercial sources (SigmaAldrich, AAT Bioquest, TCI, Method Table 2).

*Fluorophore concentrations*

The concentration of commercially available dyes was determined by their weight on a fine-balance (Sartorius Entris II) and using the molecular masses (Method Table 2). The concentration of all dyes was verified by using the Lambert-Beer law (Equation 1) and the molecular extinction coefficient according to commercial sources or literature reference (Method Table 1 and Method Table 2). All dyes were maintained in high-concentration (>1 mM) dry DMSO-$d_6$ (Carl Roth) stocks and diluted to keep the final DMSO concentration below 0.1% for *in vitro* assays and cellular experiments.

$$Abs = \varepsilon \cdot d \cdot c \quad (1)$$

*Abs – absorbance [AU], ε – molar extinction coefficient ($M^{-1}cm^{-1}$), d – optical path length (cm), c – molar concentration (M).*

**Method Table 1 | HRMS validation and extinction coefficient for custom-synthesizes dyes used in this study.**

| Dye | Chemical formular | $[M+H]^{+}_{calc}$ | $[M+H]^{+}_{det}$ | ε [$M^{-1}\cdot cm^{-1}$] |
|---|---|---|---|---|
| TMR | $C_{24}H_{22}N_2O_3$ | 387.1703 | 387.1702 | 89,000[a] |
| $JF_{549}$ | $C_{26}H_{30}N_2O_2Si$ | 429.1993 | 429.1982 | 101,00[a] |
| MaP555 | $C_{26}H_{29}N_4O_4S$ | 493.1904 | 493.1898 | 88,000[a] |
| SiR | $C_{26}H_{22}N_2O_3$ | 411.1703 | 411.1702 | 120,000[b] |
| $JF_{646}$ | $C_{28}H_{28}N_2O_2Si$ | 453.1993 | 453.1994 | 106,000[b] |

*[a] In PBS pH 7.4. [b] in activity buffer + 0.1% SDS*

**Method Table 2 | Properties of commercially available dyes used in this study.**

| Dye | source | CAS | MW [g/mol] | Ext. coeff. ε |
|---|---|---|---|---|
| Fluorescein | TCI | 2321-07-5 | 332.31 | 80,000[a] |
| R110 · HCl | Sigma-Aldrich | 13558-31-1 | 330.34 | 80,000[b] |
| R19 | TCI | 25152-49-2 | 414.50 | 90,000[c] |
| RB | TCI | 81-88-9 | 442.56 | 106,000[b] |
| R101 | Sigma-Aldrich | 116450-56-7 | 490.60 | 139,000[d] |
| PyrY | Sigma-Aldrich | 92-32-0 | 267.35 | 48,510[b] |
| PyrB | Sigma-Aldrich | 2150-48-3 | 323.46 | 106,000[b] |

*[a] in 0.1 N NaOH, [b] in dd$H_2O$. [c] in MeOH. [d] in EtOH. Extinction coefficients according to commercial sources (AAT Bioquest, Merck, TCI).*

### 3. Molecular Biology and Biochemistry

#### *Plasmid and molecular cloning*

Proteins were expressed in *E.coli* using the pET51b(+) vector (Novagen) containing a *N*-terminal $His_{10}$-tag attaches *via* a tobacco etch virus (TEV) cleavage site.[18] Protein libraries were cloned into pJYDNG vector (Addgene, #162452) containing the appS4 /Aga2p proteins for yeast surface display. Co-expressing of eUnaG2 allowed staining with bilirubin for expression normalization.[19] pcDNA5/FRT/TO expression vector (ThermoFisher Scientific) carrying different sub-cellular markers (Method Table 3) were used for mammalian expression. Direct fusion to or co-translational expression of mEGFP by using self-cleavage peptide sequence (T2A, P2A)[20, 21] was used to identify transfected cells or for the normalization of signal intensities to the protein expression. pOG44 plasmid (ThermoFisher Scientific) was used for the generation of stable cell lines. Mosaic expression in neurons of zebrafish larvae was achieved by using pTol2-*elavl3* vectors, the transgene was flanked by Tol2 sides.[22]

Gibson assembly[23] was used for molecular cloning. Site-directed mutagenesis was performed using the Q5 site-directed mutagenesis kit (NEB) according to the manufacture's protocol. Sequences were verified by Sanger sequencing (Microsynth).

**Method Table 3 | Localization sequences for mammalian cells used in this study**

| name | structure | source | name | structure | source |
|---|---|---|---|---|---|
| H2B | histone / nucleus | #169329 | LAMP1 | lysosomes | #175541 |
| $NLS_3$ | nucleus | #226514 | VIM | intermediate filament | #226518 |
| TOM20 | OMM | #135443 | CD86 | plasma membrane | #190745 |
| Sec61β | ER | #141152 | LifeAct | actin | #193328 |

*OMM – outer mitochondrial matrix, ER – endoplasmic reticulum. Source – Addgene catalog numbers.*

#### *Library generation*

A Rho-tag0.1 synthetic deep mutational scanning library (sDMSL) was purchased from Twist Bioscience. Site-saturation mutagenesis libraries (SSML) were generated by using degenerated primers (containing NNK codons) and assembly PCR[24] using the KOD Hot-Start DNA Polymerase (Merck KGaA) for DNA amplification. Both types of libraries had <70 bp overhangs to the appS4/Aga2p linker (pJYDNg backbone) for later homologous recombination in yeast. The template DNA was removed by DpnI (NEB) treatment, the DNA libraries were purified using the QIAquick PCR purification kit (QIAGEN) and 1-5 µg DNA were precipitated by using NaOAc (30 mM) and glycogen (0.5 µg/µL ThermoFisher) in isopropanol at -20 °C overnight.

#### *Recombinant protein expression, and purification from E.coli*

For recombinant protein production, *E. coli* BL21(DE3)-pLysS (Novagen) were transformed with pET51b(+) plasmids. Cultures were grown in lysogeny broth (LB) containing ampicillin ($LB^{Amp}$) at 37 °C and protein expression was induced by the addition of 0.5 mM isopropyl β-D-thiogalactopyranoside (IPTG), followed by overnight incubation at 16 °C. Cells were harvested by centrifugation (4,000 × g, 15 min, 4 °C), resuspended in His extraction buffer supplemented with 1 mM PMSF and 0.25 mg/mL lysozyme, and lysed on ice by sonication (50% duty cycle, 70% power, 7 min). The lysate was cleared by ultracentrifugation (50,000 × g, 25 min, 4 °C). Proteins were purified via immobilized metal affinity chromatography (IMAC, HisTrap FF crude columns, Cytiva) on an ÄktaPure FPLC (Cytiva), followed by buffer exchange (HiPrep 26/10 desalting colum, Cytiva) to 50 mM HEPES, 50 mM NaCl, pH 7.3 (activity buffer). For buffer composition, sees Method Table 6. The protein samples were concentrated using 10 kDa MWCO Amicon® Ultra-15 centrifugal filters. Concentration was determined by 280 nm absorbance (NanoDrop2000c, ThermoFisher Scientific) and applying the Lambert-Beer law (Equation 1) with the extinction coefficients at 280 nm according to the protein's primary sequence. Purity and molecular weight were verified by SDS-PAGE and high-resolution mass spectrometry (HRMS). Purified proteins were flash-frozen in liquid nitrogen and stored at –75 °C.

*Protein crystallization, X-ray diffraction data collection and protein structure analysis*

For protein crystallization, the $His_{10}$-TEV tag from purified protein samples was cleaved using TEV protease (weight ratio 1:30) in TEV-cleavage buffer overnight at 30 °C. The samples were filtered, purified by reverse IMAC purification (HisTrap FF crude column, Cytiva) and size exclusion chromatography (HiLoad 26/600 Superdex 75 pg column, Cytiva) both on an ÄktaPure FPLC (Cytiva). Concentrated protein samples were labeled with TMR (2x excess) for 4 h at room temperature, concentrated to 10 – 15 mg/mL and quantified using the protein extinction coefficient at 280 nm which was corrected for TMR absorbance ($TMR_{CF,\,280nm}$ = 0.16).

Protein crystals were grown at 20 °C using the vapor-diffusion method. Crystals of $CTR107^{N138A}$ / TMR were grown by mixing equal volumes of protein / TMR complex solution with reservoir solution containing 0.1 M sodium acetate pH 4.5, 0.88 M sodium dihydrogen phosphate and 1.32 M potassium hydrogen phosphate. Crystals of Rho-tag / TMR were obtained by mixing TMR complex with 0.1 M sodium phosphate citrate pH 4.2, 1.6 M sodium dihydrogen phosphate, 0.4 M potassium hydrogen phosphate. Afterwards, the crystals were gently washed with the reservoir solution containing glycerol (20% v/v total concentration) and flash-cooled in liquid $N_2$.

X-ray diffraction data were collected on the X10SA beamline at the SLS (PSI, Villigen, Switzerland) at 100 K. Diffraction data were processed with XDS[25]. The structure of $CTR107^{N138A}$ / TMR was determined by molecular replacement (MR) using Phaser[26] and the CTR107 coordinates (PDB ID 5KAX) as a search model. The structure of Rho-tag/TMR was determined using $CTR107^{N138A}$ / TMR model. Twinning was indicated for Rho-tag/TMR data and the *k, h, -l* twin operator was used during refinement. Grade server[27] was used to produce geometrical restraints for the TMR ligand. The final models were optimized in iterative cycles of manual rebuilding using Coot [28] and refinement using Refmac5 [29] and phenix.refine [30]. Data collection and refinement statistics are summarized in Supplementary Table 8, model quality was validated with MolProbity [31] as implemented in PHENIX. Atomic coordinates and structure factors have been deposited in the Protein Data Bank under accession codes: 9RTL ($CTR107^{N138A}$ / TMR), 9RTM (Rho-tag / TMR). Crystal structure were visualized and analyzed with PyMOL 2.6.[32]

*Yeast culture and transformation*

The *S. cerevisiae* strain EBY100 (ATCC) was cultured at 30 °C, 250 rpm in YPD or SDCAA media using sterile Erlenmeyer flasks or 14 mL polypropylene tubes (Falcon). Cultures were passaged every 1 - 3 days with a 10-fold dilution. Electrocompetent EBY100 cells were prepared according to the following protocol. A single colony was used to inoculate a 50 mL YPD culture which was grown overnight. The next day, the culture was diluted to $OD_{600}$ 0.3 in 100 mL YPD and grown until $OD_{600}$ 1.4 was reached. Tris-DTT (800 µL) and Tris-LiAc (2 mL) were added for 15 min and incubated at 30 °C, 250 rpm. Cells were harvested (2,500 × g, 3 min, 4 °C), washed once with 25 mL cold electroporation buffer, and centrifuged again. Pellets were resuspended in 300 µL electroporation buffer, aliquoted (12 x 50 µL) on ice, and used immediately for electroporation with 1–5 µg DNA precipitated DNA. Cells were transferred to pre-chilled 0.2 cm gap electroporation cuvettes (Bio-Rad), and electroporated using a Gene Pulser Xcell (0.54 kV, 25 µF, infinite resistance, exponential decay). Cells were recovered in 2 mL YPDS at 30 °C, shaking for 1 h and afterwards expanded in 100 mL SDCAA at 30 °C for 2 days.

*YSD protein expression, staining and sorting*

SGCAA medium was used for induction and cells were grown at 30 °C, 250 rpm for 20 h. About $10^8$ cells were harvested (14,000 × g, 1 min), resuspended in 500 µL PBS with 10 µM bilirubin, and incubated at 8 °C on a rotating wheel for 30 min. Cells were washed twice with PBS (500 µL each) and resuspended in 500 µL PBS. For labeling, they were mixed 1:1 with 2× dye solution (final concentration 500–2 nM). Samples were filtered through 5 mL round-bottom tubes with cell strainer caps (Falcon) and analyzed directly by FACS.

Cell enrichment was performed on a BD FACS Melody sorter (100 µm nozzle, FSC ND 1.5 filter). PMT voltages were set based on negative controls (no protein expression). Gates were applied for FSC, SSC, and fluorescence; signals $>10^3$ (bilirubin or rhodamine dyes) were considered positive, background $\leq 10^2$. Filter configurations are collected in Method Table 4. In the initial round, at least twice the theoretical library size was sorted; subsequent rounds collected 10,000 cells each. Following transformation or sorting, Penicillin-Streptomycin (50 U/mL, Gibco) was added to the SDCAA medium.

*Plasmid isolation and NGS analysis*

The plasmid DNA from each sorting round was extracted using the Zymoprep Yeast Plasmid Miniprep II kit (Zymo Research) according to the manufacturer's suggestions. The randomized regions were amplified *via* KOD PCR reaction using primers containing unique barcodes (NGS adaptor ligation oligos, Eurofins Genomics) analyzed by NGS sequencing (Illumina technology, Eurofins Genomics). The NGS results were analyzed using a custom-build pipeline as described previously.[33] Alternatively, plasmid DNA was transformation into *E.cloni* 10G and single colonies were analyzed by Sanger sequencing (Eurofins).

**Method Table 4 | Filter settings used at FACS Melody for YSD screens**

| protein | Excitation laser | Channel | Emission filter |
|---|---|---|---|
| Bilirubin | 488 nm | FITC | 527/32 |
| MaP555 | 561 nm | PE | 582/15 |
| $JF_{646}$ | 640 nm | APC | 660/10 |

### 4. Biochemical and Spectral Characterization

*Affinity measurement*

Serial protein dilutions (0 – 200 µM) were prepared in activity buffer and transferred to black flat bottom low-volume 384-well plates (Greiner). Rhodamine dyes (0.5 – 20 nM) were added in activity buffer containing 1% (w/v) bovine serum albumin (BSA, Fraction V, Roth) using a 384-channel VIAFLO pipettor (Integra) at a 1:1 mixing ratio. Fluorescence intensity (FI, SiR / $JF_{646}$) or fluorescence polarization (FP, all other dyes) measurements were conducted on a Spark20M microplate reader (Tecan) at 25 °C in technical triplicates. Excitation and emission settings are summarized in Method Table 5.

For data processing, technical triplicates were averaged and normalized between 0 (*i.e.* free dye) and 1 (fully bound dye). The dissociation constants $K_D$ were determined by fitting a single-site binding model (Equation 2) to the data. Results are reported as mean ± standard deviation from at least three independent experiments (N ≥ 3), unless otherwise mentioned. To compare between dyes or proteins, $\log_{10}$ fold changes relative to a reference *e.g.,* parental protein were calculated.

$$Y = A + \frac{(B-A)^h}{1+\left(\frac{K_D}{[P]}\right)^h} \quad (2)$$

*Y: fluorescence intensity [a.u.] or fluorescence polarization [mFP], A: min. FI / FP, B: max. FI / FP, [P]: protein concentration [mol/L], h: Hill coefficient, $K_D$: dissociation constant [mol/L].*

**Method Table 5 | Excitation and emission settings used for affinity measurement**

| Fluorophore | $\lambda_{ext}$ [nm] | $\lambda_{em}$ [nm] | Gain set [%] | method |
|---|---|---|---|---|
| Fluorescein | 470 ± 20 | 520 ± 20 | 80 | monochromator |
| R110 / R19 | 510 ± 20 | 560 ± 20 | 80 | monochromator |
| RB / R101 / PyrY / PyrB | 535 ± 12.5 | 595 ± 17.5 | 80 | filter |
| TMR / $JF_{549}$ | 535 ± 12.5 | 595 ± 17.5 | 80 | filter |
| SiR / $JF_{646}$ | 620 ± 10 nm | 665 ± 10 nm | 100 | monochromator |

*Stopped-flow kinetic fluorescence anisotropy*

Binding kinetics were recorded on a BioLogic SFM-400 stopped-flow system (BioLogic Science Instruments). Equal volumes of Rho-tag (4 µM) and rhodamine dye (1 µM) in activity buffer were rapidly mixed (singlemix sequence). Fluorescence anisotropy changes were monitored over 300 s at 0.25 ms intervals. Background signal was assessed by mixing dye with 1% (w/v) BSA. The instrument-determined dead time (3.7 ms) was subtracted from all time points. Measurements were performed at 37 °C. TMR was excited at 553 nm and recorded using a 570 nm emission long pass filter, SiR / $JF_{646}$ were excited at 650 nm and recorded using a 665 nm emission long pass filter. Each condition was recorded in 12 technical replicates, averaged, background-corrected, and normalized to the highest observed anisotropy (B). The association rate constant ($k_{on}$) was obtained by fitting data to a second-order integrated rate equation to the data (Equation 3).

$$\mathrm{FA} = \mathrm{B} + \frac{\left(0-\frac{B}{[D]}\right) \cdot \left([D] \cdot ([D]-[P])\right) \cdot e^{([D]-[HT]) \cdot k_1 \cdot t}}{D \cdot e^{(([D]-[P]) \cdot k_1 \cdot t)-[HT]}} \tag{3}$$

*FA: fluorescence anisotropy, B: max. FA, [D]: dye concentration, [P]: protein concentration, t: time [s],* $k_1$*: on-rate* [$M^{-1}$ $s^{-1}$]*.*

Together with experimentally evaluated $K_D$-values, the unbinding kinetic constants were calculated using Equation (4).

$$K_D = \frac{k_{off}}{k_{on}} \leftrightarrow k_{off} = K_D \cdot k_{on} \tag{4}$$

*$K_D$ – dissociation constant [M], $k_{off}$ – unbinding rate constant [$s^{-1}$], $k_{on}$ – binding rate constant [$M^{-1}s^{-1}$]*

*nanoDSF*

The thermal stability of Rho-tag variants was assessed using nano differential scanning fluorimetry (nanoDSF) on a Prometheus NT48 (NanoTemper Technologies). Protein samples (100 µM in activity buffer) were subjected to a thermal gradient from 20° C to 95° C at 1 °C / min. The intrinsic fluorescence ratio (350 / 330 nm) of tryptophan / tyrosine residues was monitored, and the melting temperature ($T_M$) was determined as the inflection point of this ratio curve. Each sample was analyzed in technical duplicate (N=2), and results are reported as mean ± standard deviation.

*Fluorescence emission spectra*

To evaluate fluorescence turn-on or quenching ($F_I/F_{I0}$), rhodamine dyes (2 – 50 nM) were incubated with or without excess Rho-tag (20 µM) in activity buffer containing 0.5% BSA (w/v). Measurements were performed in black 384-well plates (Greiner) using the Spark20M microplate reader (Tecan) in technical replicates. TMR was excited at 530 nm and the emission was recorded from 556 – 706 nm. SiR was excited at 626 nm and the emission was recorded from 656 – 800 nm. $F_I/F_{I0}$ was defined as the ratio of fluorescence intensity with protein to that without at the emission maximum.

*Extinction coefficient*

Serially fluorophore dilutions (0 – 3 µM) were prepared in activity buffer in presence or absence of 20 µM Rho-tag. Measurements were performed in 200 µL volumes in clear-bottom non-binding 96-well plates (Greiner). Absorbance at $\lambda_{max}$ (TMR / $JF_{549}$ – 555 nm, SiR / $JF_{646}$ – 646 nm) was recorded on the Spark20M microplate reader (Tecan). Following baseline correction, the absorbance was plotted against the dye concentration and the data were fitted linearly. Path length was calibrated using the following reference extinction coefficients: TMR in PBS ε 89,000 $M^{-1}\cdot cm^{-1}$[16] and SiR in 0.1% SDS in activity buffer ε 120,000 $M^{-1}\cdot cm^{-1}$[15]. The extinction coefficients were calculated *via* the Lambert-Beer law (Equation 1).

*Quantum yield*

Absolute quantum yields were determined at room temperature using a Hamamatsu Quantaurus-QY spectrometer (C11347 model). Rhodamine dyes (1 µM) were tested with and without excess Rho-tag (20 µM) in activity buffer + 0.5% BSA. Samples were prepared in 800 µL glass vials (TMR, $JF_{549}$) or 2 mL quartz cuvettes (SiR, $JF_{646}$) and measured at $\lambda_{abs}$ (TMR – 555 nm, $JF_{549}$ 550 nm, SiR / $JF_{646}$ – 646 nm). Quantum yield represent as mean ± standard deviation from three technical replicates. The brightness was thereafter calculated using Equation 5.

$$\text{brightness} = \varepsilon \cdot \Phi \qquad (5)$$

*ε - extinction coefficient [$M^{-1}cm^{-1}$], Φ [0 – 1]*

*Bioluminescence scans and BRET ratio*

Bioluminescence was measured in white non-binding 96-well plates (PerkinElmer, 200 µL volume) on the Spark20M (Tecan) with a luminescence module. Assay mixture contained 2 nM NanoLuc fusion protein, 50 nM TMR, Nano-Glo® substrate (Promega 1:1000) in activity buffer + 0.2 mg/mL BSA. Emission spectra (398–653 nm) were collected with 15 nm steps and 1000 ms integration. The BRET ratio was defined as $I_{593}/I_{459}$.

**Method Table 6 | Buffer and media composition used for biochemical experiments in this study**

| Buffer | Composition |
|---|---|
| $LB^{Amp}$ | 5 g/L yeast extract, 10 g/L peptone, 0.1 g/L ampicillin |
| Activity buffer | 50 mM HEPES, 50 mM NaCl, pH 7.3 |
| Extraction buffer | 50 mM $KH_2PO_4$, 300 mM NaCl, 5 mM imidazole, pH 8.0 |
| IMAC wash buffer | 50 mM $KH_2PO_4$, 300 mM NaCl, 10 mM imidazole, pH 7.5 |
| IMAC elution buffer | 50 mM $KH_2PO_4$, 300 mM NaCl, 500 mM imidazole, pH 7.5 |
| TEV-cleavage buffer | 25 mM $Na_2HPO_4$, 200 mM NaCl, pH 8.0 |
| YPD medium | 20 g/L glucose, 20 g/L peptone, 10 g/L yeast extract, autoclaved (prior to addition of glucose) |
| SDCAA drop out medium | 20 g/L glucose, 6.7 g/L yeast nitrogen base (Difco), 5 g/L Bacto casamino acids (without tryptophan), 38 mM |
| SGCAA drop out medium | 20 g/L galactose, 6.7 g/L yeast nitrogen base (Difco), 5 g/L Bacto casamino acids (without tryptophan), 38 mM $Na_2HPO_4$ × 12 H2O, 62 mM $NaH_2PO_4$ × H2O, autoclaved (prior to addition of galactose) |
| YPD agar plates | YPD medium, 15 g/L agar (1.5 % w/v) |
| SDCAA drop out agar plates | SDCAA drop out medium, 3 g/L agar, 1 M sorbitol |
| Tris buffer | 1 M Tris-HCl, pH = 8 |
| Tris-DTT buffer | 0.39 g DTT (2.5 M) dissolved in Tris buffer (800 µL), filtered |
| Tris-LiAc buffer | 1.02 g LiAc × 2 H2O dissolved in Tris buffer (2 mL), filtered |
| Electroporation buffer | 10 mM Tris-base, 270 mM Sucrose, 2.1 mM $MgCl_2$ × 6 $H_2O$, pH = 7.5, autoclaved |
| YPDS | 1:1 mixture of YPD and 1 M sorbitol |

*Buffers/media were prepared in Milli-Q® water*

### 5. Cell Biology and Microscopy

*Mammalian Cell-culture*

U-2 OS Flp-In T-Rex cells[34] (referred to as U2OS), and Hela Kyoto Flp-In cells[35] (referred to as HeLa cells), were cultured in T-25 cell culture flasks (Greiner) and maintained in high-glucose Dulbecco's Modified Eagle Medium (DMEM GlutaMAX™, phenol-red, Gibco) supplemented with 10% (v/v) fetal bovine serum (FBS). The cells were stored in a humidified incubator at 37 °C with 5% $CO_2$ and routinely passaged every 2–3 days using phosphate-buffered saline (PBS, pH 7.4, Gibco) and TrypLE™ Select Enzyme (1x, phenol-red free, Gibco).

Stable U2OS cells lines were established using the Flp-IN T-REx™ system (ThermoFisher Scientific) according to previous protocols[17] and eventually sorted for transgene expression using FACS on the a BD FACS Melody sorter. Unless otherwise state, microscopy experiments within this study were performed using stable cell-lines. Recombinant adeno-associated viruses (rAAVs) carrying TOM20-HaloTag7 transgene were produced as described previously.[17]

*Cell seeding, fixation and staining*

Two days before imaging experiments, between 80,000 and 300,000 cells per well were plated in either tissue culture-treated 96-well µ-Plates (ibidi, square glass bottom) or 8-well µ-Slides (ibidi) having optically clear glass bottoms. Unless otherwise mentioned, stable cell lines were co-seeded with 20% wild-type U2OS cells (lacking protein tag expression). The following day and at least 16 hours prior to imaging or FACS, transgene expression was induced by adding 0.1 - 0.5 µg/mL doxycycline (Sigma-Aldrich). For transient transfection, 100 ng plasmid DNA per well was delivered using Lipofectamine 3000® reagent (ThermoFisher Scientific) according to the manufacturer's protocol (overnight incubation).

Cells were rinsed with PBS and fixed using 4% (v/v) paraformaldehyde (Electron Microscopy Sciences) in PBS pre-warmed to 37 °C for 20 minutes. Residual fixative was neutralized with 50 mM ammonium chloride (Sigma-Aldrich) in PBS, after which cells were washed twice with PBS. Fixed samples were stored at 4 °C in PBS containing 0.5% BSA, for up to one week.

**Fluorophores (5–500 nM) were diluted in imaging medium (composition sees**

Method Table 7) for live cell or pre-fixation labeling or in PBS containing 0.5% BSA for post-fixation labeling. For HaloTag7 labeling, the ligands were incubated for at least 30 minutes at 37 °C on the cells and washed as indicated before imaging.

*Zebrafish husbandry and embryo handling*

Zebrafish (*Danio rerio*) larvae were prepared utilizing the infrastructure provided by Prof. Jochen Wittbrodt (COS Heidelberg). Parental zebrafish stocks were maintained (fish husbandry, permit number 35-9185.64-BH to Jochen Wittbrodt) in accordance with local animal welfare standards (Tierschutzgesetz §11, Abs. 1, Nr. 1) and with European Union animal welfare guidelines. The fish facility is under the supervision of the local representative of the animal welfare agency. The herein described experiments on zebrafish larvae, followed by their euthanization, were performed before 5 *dpf*.

The animals were kept under a controlled 14:10 hours light:dark cycle and in a constant recirculating system at 28 °C, with water conditions maintained at pH of 7-7.5, and a conductivity of 600 µS.[36] Embryos obtained through natural mating were raised in zebrafish medium, with medium exchanged every other day. Developmental staging was recorded as days post-fertilization (dpf), based on standard incubation at 28.5 °C. All embryos used in this work carried the Casper genetic background, having the mitfa$^{-/-}$[37] and mpv17$^{-/-}$[38] mutations and melanophore and iridophore pigmentation deficiency for improved optical accessibility.[39]

Wild-type *Casper* zebrafish eggs were injected at the single-cell stage (0 *dpf*) with *in vitro* transcribed mRNA encoding Tol2 transposase (15 ng/µL), alongside with a pTol2 plasmid DNA containing the *elavl3:Rho-tag-/SiR-tag/HaloTag7-P$_{30}$-mEGFP-NLS$_3$* constructs (10 ng/µL). Transgene expression was driven by a truncated pan-neuronal *elavl3* promoter (mosaic expression). At 3 *dpf*, hatched embryos exhibiting mEGFP fluorescence were selected under an epifluorescence microscope (excitation: 488 nm; emission: 530/30 nm). For imaging experiments, larvae (3–4 *dpf*) were embedded in 1.3% low-melting-point agarose supplemented with 1x tricaine (0.168 mg/mL) to anesthetized the larvae. The embryos were centrally placed within glass-bottom 100 mm × 35 mm plastic dishes (4 mL, MatTek glass, 10 mm Glass Diameter, uncoated). The fluorophores were added to a final concentration of 250 nM and the labeling was followed under a confocal microscope (SP8, Leica). To test if the fluorescent label can be removed, fully labeled larvae were release from the agarose, rinsed twice, left freely swimming in fresh medium for 10 min and mounted again for imaging.

To display the full brain volume, maximal projections of 200 µm optical sections were created using Fiji. Co-localization between the fluorescent probe and the mEGFP expression marker was probed using the Fiji plug-in 'Coloc 2' by their $R^2$ values.

**Method Table 7 | Buffer and media composition used for cell-biology in this study**

| Buffer | Composition |
|---|---|
| Cell growth medium | DMEM phenol-red GlutaMAX™ with 4.5 g/L glucose, pyruvate (1 x), 10 % FBS (Gibco) |
| Cell imaging medium | DMEM phenol-red free with 4.5 g/L glucose (1 x), GlutaMAX™ (1x), sodium pyruvate (1x), HEPES (50 mM), 10 % FBS (all Gibco) |
| Zebrafish medium | 0.3 g red sea salt in 1 L VE water |

*Confocal Fluorescence Microscopy*

Confocal imaging was carried out on either a Leica DMi8 inverted microscope equipped with a TCS SP8 X scanhead and a 40×/1.10 motCORR water immersion objective or a Stellaris 5 inverted microscope equipped with a 20×/0.75 dry objective (both Leica Microsystems). The correction ring on the water objective was adjusted via reflection-based alignment in FCS mode. SuperK white light lasers (WLL, 488 - 650 nm) were used to excite the fluorophores. Fluorescence signals were captured using hybrid detectors (HyD). Dual-color imaging was done sequentially to prevent bleed-through. Images were recorded at 400 - 600 Hz scan speed, with up to 3× line accumulation, zoom factors of 0.75–3.0, and image dimensions of 1024×1024 or 2048×2048 pixels at 12- or 16-bit depth. Unless noted otherwise, a pinhole of 1 Airy unit was used. Further acquisition settings are summarized in Method Table 8. Z-stacks were acquired using steps of 1 or 2 µm. During live imaging, mammalian cells were kept at 35 °C with 5% $CO_2$; imaging of zebrafish was conducted at 30 °C.

*Background staining*

To assess non-specific binding, wild-type U2OS cells were exposed to 200 nM fluorophore at 37 °C for 1 hour. The laser was adjusted to saturate the brightest signal before acquiring confocal images.

*Washing protocols*

U2OS cells expressing H2B-Rho-tag-T2A-mEGFP were labeled with 50 nM TMR. Three washing strategies were evaluated to remove residual fluorophore: *i)* single wash with 200 µL pre-warmed imaging medium; *ii)* two times medium replacement; iii) incubation with 5 µM Rho-tag for 5 min followed by medium replacement with fresh, pre-warmed imaging medium.

*Live-cell kinetics*

U2OS cells stably expressing $P_{30}$-mEGFP-$NLS_3$ fused to Rho-tag or HaloTag7 were incubated with 50 nM TMR or TMR-HTL, respectively. The laser settings were adjusted to the signal after full labeling (1 h at 37 °C). Baseline images were collected prior to dye addition. Subsequently, the fluorophore was added in a 1:1 dilution and image were acquired with a dead-time of 12 sec. Confocal z-stacks were recorded with a frame-rate of ~25 sec per stack. Live-cell kinetic curves were generated by extracting the mEGFP-normalized nuclear signal over time.

*Photobleaching*

Similar to live-cell kinetic experiments, fully labeled cells (50 nM fluorophore, 1 h at 37 °C) were subjected to photobleaching using the Stellaris5 FRAP module. The bleaching protocol employed iterative image acquisition (1% laser power) and bleaching (100% laser power, 555 nm, 5 times repetition) in circular ROIs. Settings include *zoom-in* mode, *zero background* enabled, and auto-deletion of bleached frames. This sequence was repeated 30 times consecutively, generating one image every 90 seconds. Bleaching curves were generated from at least 25 nuclear signals (N=25) from at least 3 images by plotting the nuclear signal over time normalized to the initial frame.

*Image processing and cellular signal quantification*

Images were processed using Fiji software (v1.54p)[40]: brightness and contrast were adjusted, maximum intensity projections generated, and 'Hot' LUTs applied for pixel intensity visualization. Gaussian blur and scale bars were added for image representation. Fluorescence signals under different staining conditions were compared by their expression-normalized signal intensity ($I_{norm}$) as defined in Equation (6). Nuclear fluorescence ($I_{nuc}$ for H2B-POI-T2A-mEGFP or POI-$P_{30}$-mEGFP-$NLS_3$ constructrs) and cytosolic GFP signal ($I_{GFP}$) were measured using circular ROIs across at least three fields of view, analyzing ≥60 cells. Background signal ($I_{bg}$) was determined from co-plated wild-type U2OS cells and subtracted from all measurements.

$$I_{norm} = \frac{(I_{nuc} - I_{bg})}{I_{GFP}} \tag{6}$$

*$I_{norm}$ - expression normalized fluorescence signal $I_{nuc}$ – nuclear signal intensity from cells expressing H2B-Rho-tag or -HaloTag7. $I_{bg}$ – nuclear signal intensity from wild-type cells. $I_{GFP}$ – cytosolic GFP signal (expression level).*

The signal-over-background (S/B) was determined as the ratio of nuclear signal from cells expressing a Rho-tag variants or HaloTag7 (signal) to the nuclear signal from non-expressing cells (background) according to Equation (7).

$$S/B = \frac{I_{nuc}}{I_{bg}} \tag{7}$$

*S/B – signal-over-background, $I_{nuc}$ – nuclear signal intensity from cells expressing H2B-Rho-tag variants or -HaloTag7. $I_{bg}$ – nuclear signal intensity from wild-type cells. $I_{GFP}$ – cytosolic signal GFP signal (expression).*

*Live-cell STED microscopy*

STED imaging was performed on an Abberior STED Expert Line 595/775/RESOLFT QUAD scanning microscope equipped with a UPlanSApo 100x/1.4 oil immersion objective lens, a 640 nm excitation laser and a 775 nm STED laser (Abberior Instruments GmbH). Fluorescence was detected on avalanche photodiodes (APD) from 650 – 757nm using spectral detection. Images were acquired in 60 x 60 µm (overview) 10 x 10 µm (magnification) dimensions. Standard settings include a pixel dwell time of 10 µs and a pixel size of 80 nm (overview) or 30 nm (magnification) with 5 average line scans. For data representation, 'Hot' LUT were applied according to the reference bars.

*Determination of the single-molecule binding kinetics and SMLM Imaging*

For single-molecule kinetic measurements, 8-well chamber slides (Sarstedt) were coated with poly-L-lysin-grafted polyethylene glycol functionalized with a peptide (CGRGDS, PLL-PEG-RGD) for cell adhesion using a protocol published previously.[41] To synthesize PLL-PEG-RGD, maleimide-PEG-NHS was reacted with Ac-CGRGDS-$NH_2$ in HBS buffer (pH 7.5) with 1 mM EDTA, then reacted with PLL in HBS buffer (pH 8.0) overnight. Chamber slides were subjected to plasma cleaning (Zepto B) with $N_2$ for 15 min and 25 µL of 0.8 mg/mL PLL-PEG-RGD was added to each well and incubated of 1 h at 37 °C. After incubation, 8-well chamber slides were dried under a sterile bench. For fixed and live-cell SMLM imaging, 8-well chamber slides (Sarstedt) were coated with fibronectin (F0895, 15 µg/mL, Merck) diluted in DPBS (Thermo Scientific) for 1 h at 37 °C, after incubation, the chamber slides were dried under a sterile bench for 30 min. U2OS cells were seeded to a density of $10^4$ cells/well into the coated chamber slides and protein expression was induced by adding 100 ng/mL doxycycline (Sigma Aldrich).

Cells were fixed with pre-warmed 4% formaldehyde (FA, Thermo Scientific) and 0.1 % glutaldehyde (GA, Electron Microscopy Science) in 1x DPBS for 20 min at room temperature. Gold beads (90 or 100 nm diameter, Nanopartz) were diluted 1:5 in DPBS, added to the cells, and incubated for 20 min at room temperature. The cells were labeled with TMR or SiR in DPBS (fixed) or FluoroBrite™ DMEM (Gibco) medium (live). 3 nM dye were used to determine the binding kinetics, 1 nM for fixed-cell SMLM and 5 – 10 nM for live-cell SMLM.

SMLM imaging was performed on a N-STORM microscope (Nikon) equipped with an 100x/1.49 oil immersion objective (Apo TIRF) and an EMCCD camera (DU-897U-CS0-#BV, Andor Technology). A 561 nm laser with an intensity range of 10.4 kW/cm$^2$ was used to excite TMR and a 647 nm laser with an intensity range of 8.34 kW/cm$^2$ was used to excite SiR under total internal reflection fluorescence (TIRF) mode. LCControl (Agilent) and NIS Elements (Nikon) software were used for setup control. The data was acquired using MicroManager (v1.4.22)[42, 43]. For fixed-cell SMLM, 150 ms exposure time, 200 EM gain, 3x preamp gain, and 17 MHz readout rate with activated frame transfer were used to record 25,000 frames per cell. For live-cell SMLM, cells were maintained a constant temperature of 37 °C in a stage top incubator (Okolab, Naples, Italy). Data was recorded with 50 ms exposure time, 50 EM gain, 1x preamp gain, and 5 MHz readout rate with activated frame transfer. The 561 nm laser was used at an intensity of 8.83 kW/cm$^2$ in highly inclined and laminated optical sheet (HILO) mode. 10,000 frames were recorded to generate a live-cell SMLM videos.

*SMLM data analysis*

Fixed-cell SMLM data were analyzed using Picasso (v0.7.0.)[44]. Single molecules in each frame were identified using the *localize* module with maximum likelihood estimation for integrated Gaussian parameters. After localization, the lateral drift was corrected in the *render* module using gold beads as fiducial markers, followed by filtering of signals from out-of-focus planes using the *filter* module by applying a range of 0.5-1.5 for the width and height of the point-spread function (sx, sy), 0 - 0.25 for the localization precision (lpx, lpy) and 0 - 0.25 for the ellipticity. Localizations presumably originating from the same fluorophore in consecutive frames were linked in a radius of 6x NeNA (maximum 0.3-pixel size). For further kinetic analysis of CD86 tagged with Rho-tag or SiR-tag, DBSCAN was performed with 2x NeNA and a minimum density of 7. Clusters are filtered for their mean frame, standard deviation of the mean frame, and cluster area using the *clusterfilter* function of Picasso or custom python script. For mean frame and standard deviation of the mean frame, 95% of clusters are kept. For cluster area, clusters with an area bigger than 0.25 px$^2$ were filtered out. Then, the bright time $\tau_b$ from the filtered DBSCAN file was fitted to a single exponential function (Equation 8) using OriginPro (version 2019).

$$P(t) = \left(1 - exp\left(\frac{t}{\tau_b}\right)\right) * a + b \tag{8}$$

*Data analysis of live-cell SMLM*

Live-cell SMLM data were analyzed as previously described.[45] DeepSTORM 2D neural network was trained using simulated data and used to predict live-cell high-density SMLM data of Sec61β-Rho-tag. After the prediction, a live-cell movie was generated using a custom-written Python script based on the video generation script published at https://github.com/alonsaguy/DBlink.[46] Using this script, super-resolution images were predicted from batches of 200 frames of the high-density data set using a temporally moving window with a 180 frame overlap between frame batches.

*MINFLUX Microscopy*

For MINFLUX imaging, U2OS cells stably expressing VIM-Rho-tag were seeded onto 18 mm diameter glass coverslips (Carl Roth) and cultured to approximately 60% confluence. Cells were chemically fixed following the procedure described above. Each of the following step included three washes with PBS containing 1% (w/v) BSA. Cells were permeabilized with 0.1% Triton X-100 for 10 min, followed by incubation for 1 h at room temperature with a primary mouse anti-vimentin antibody (Sigma-Aldrich, cat. V6389, 1:500), followed by 1 h incubation with a Alexa Fluor 647-conjugated goat anti-mouse secondary antibody (Thermo Fisher, cat. A32728, 1:1000) at room temperature. After antibody labeling, cells were incubated for 5 min at room temperature with fiducial markers (NanoParz, cat. A12-40-980-CTAB-DIH1-25) and stained using 0.2 nM TMR in PBS containing 1% (w/v) BSA.

MINFLUX imaging was performed using a commercial 3D MINFLUX system (Abberior Instruments) mounted on a motorized Olympus IX83 inverted microscope and armed with a 561 nm MINFLUX laser. Fluorescence signals were collected on avalanche photodiodes (APDs) within the 580 – 630 nm detection window. The pinhole was set to 0.67 Airy units, and the initial periscope excitation power was approximately 35 µW. Data were acquired for 2 h using a default

2D imaging sequence, with a localization range L reaching 40 nm and limited to 100 photons in the last iteration step.
For visualization, ‘Hot’ lookup tables were applied. The localization precision was calculated and the trace iD over time (TiM) was plotted using the PARAFlux software (v1.22, copyright Tobias Weihns, Abberior Instruments GmbH).

*Single particle tracking*
For single-particle tracking, coverslips (25 mm diameter, 0.17 mm thickness, VWR) were cleaned by sonicating in isopropanol for 20 min and plasma-cleaning for 10 min as described above. Coverslips were coated with poly-L-lysin-peptide by applying 8 µL of 0.8 mg/mL PLL-PEG-RGD (preparation described above) between two coverslips and incubation for 1.5 h at room temperature. After incubation, $ddH_2O$ was added to separate the coverslips before drying and placing them in 6-well plates. Cells were seeded at a density of 4 x $10^4$ cells/well and grown for 3 days at 37 °C with 5% $CO_2$ and protein expression was induced with 100 ng/mL doxycycline in growth medium.
Prior to SPT experiments, the coverslips were placed in a home-built coverslip holder and TMR or SiR were added in FluoroBrite™ DMEM medium to a final concentration of 1 nM. The samples were equilibrated to room temperature for 15 min. SPT experiments were performed on a N-STORM microscope (Nikon) at a constant temperature of 25 °C. The 561 nm laser was used at 0.4 kW/$cm^2$ intensity in TIRF mode using the following camera settings: 20 ms exposure time, 200 EM gain, 3x preamp gain, 17 MHz readout rate, and activated frame transfer. 1,000 frames were recorded per cell, using a frame size of 256 x 256 px and a pixel size of 157 nm. 10-30 cells per coverslip were measured within 30 min. Transmitted light images were recorded before each SPT experiment. For each coverslip, background films were recorded applying the same settings in regions without cells.  Long-term SPT experiments were conducted for 30 min (90,000 frames) using the same experimental settings.

Data were analyzed according to previously published workflows[41, 47, 48]. In brief, single emitters were localized with the Fiji plugin ThunderSTORM.[49] The trajectories of single molecules were reconstructed using swift (v0.4.3)[50] and the diffusion was analyzed with SPTAnalyser (v1.2.0)[47]. Parameters for all analysis steps were determined according to the SPTanalyser manual (v1.2.0). The following parameters were used for swift: *diffraction_limit* = 10 nm, *exp_displacement* = 114 nm, *p_bleach* = 0.059, and *p_switch* = 0.01. SPTAnalyser parameters were set to *id* = seg, $D_{min}$ = 0.0033 µm²/s, and *minimum trajectory length* = 20.

### 6. Statistical Analysis and Reproducibility

All biochemical assays were carried out in a minimum of three independent experiments ($N \geq 3$), unless otherwise stated. Data are reported as mean values with corresponding standard deviations. For variables derived from products and quotients the error was calculated using propagation of uncertainty, as described by Equation 9.

$$\sigma = |C| \sqrt{\left(\frac{\sigma_A}{A}\right)^2 + \left(\frac{\sigma_B}{B}\right)^2} \quad (9)$$

*$\sigma$: standard deviation. A, B: variables. $\sigma_A$, $\sigma_B$: standard deviation of variables A and B. $C = A \cdot B$ or $C = \frac{A}{B}$.*

Cell-based experiments were performed at least twice ($N \geq 2$), yielding consistent results across replicates. For quantitative analyses, at least 60 individual cells were evaluated, distributed across at least three individual microscopy images. For comparisons of large sample sets (≥ 60 cells), standard error of the mean (SEM) was applied as specified in figure captions. Statistical significance was determined by two-tailed Student’s t-test. For cell populations, Welch’s correction was applied using OriginLab 2021b software. Results with p-values ≥ 0.05 were classified as not significant (n.s.), whereas p-values < 0.05 (*) were considered significant, as indicated in the figure captions.

**Method Table 8 | Confocal imaging acquisition parameters.**

| Fig. | system | objective | $\lambda_{Ex}$ nm (%) | $\lambda_{Em}$ nm | size [pxl] | scan speed [Hz] / dwell time [µs] | optical zoom | lines |
|---|---|---|---|---|---|---|---|---|
| **3A** | **A** | 40x water | 488 (7)<br>555 (3) | 500 – 540<br>565 -619 | 1024 x 1024 | 600 | 0.75 | uni, 1 |
| **3D** | **A** | 40x water | 488 (7)<br>560 (2) | 500 – 540<br>570 - 670 | 1024 x 1024 | 600 | 1 | uni, 1 |
| **3F** | **C** | 60x oil | 561 (5) | 571 – 650 | 60 x 60 | 15 | - | 3 |
| **3G** | **A** | 40x water | 550 (8)<br>645 (5) | 560 -620<br>655 - 750 | 2048 x 2048 | 400 | 3 | uni, 1 |
| **7A, B** | **A** | 20x dry | 488 (10)<br>550 (10) | 500 – 530<br>560 - 610 | 1024 x 1024 | 600 | 0.75 | bi, 3 |
| **7C** | **A** | 20x dry | 488 (10)<br>550 (10) | 500 – 530<br>560 - 610 | 1024 x 1024 | 600 | 0.75 | bi, 3 |
| **7D** | **A** | 20x dry | 488 (20)<br>635 (20) | 500 - 550<br>650 - 720 | 1024 x 1024 | 600 | 0.75 | bi, 3 |
| **S3** | **A** | 40x water | 560 (2) | 570 - 670 | 1024 x 1024 | 600 | 1 | uni, 3 |
| **S5** | **A** | 40x water | 488 (7)<br>560 (2) | 500 – 540<br>570 - 670 | 1024 x 1024 | 600 | 1 | uni, 3 |
| **S8** | **B** | 20x dry | 555 (1) | 565 -665 | 1024 x 1024 | 600 | 1 | uni, 2 |
| **S9** | **A** | 20x dry | 555 (1) | 565 -665 | 1024 x 1024 | 600 | 1 | uni, 2 |
| **S10** | **A** | 40x water | 488 (20)<br>550 (2) | 500 -535<br>565 - 650 | 1024 x 1024 | 400 | 0.75 | uni, 1 |
| **S12** | **A** | 20x dry | 488 (10)<br>550 (10) | 500 – 530<br>560 - 610 | 1024 x 1024 | 600 | 0.75 | bi, 3 |

*Instrument **A**: Leica DMi8 microscope (SP8) – WLL,*
*Instrument **B**: Leica Stellaris5 - WLL*
*Instrument **C**: STED Infinite line. Excitation line: 561 nm; Emission: 571 – 650 nm.*
*Scan direction: Unidirectional (uni) or bidirectional (bi)*

### Supplementary Videos

**Video S1. Single-molecule SMLM imaging of Vimentin-Rho-tag labeled with TMR in fixed U2OS cells.**
The video represents the raw data underlying Figure 6C. Scale bar: 2 µm.

**Videos S2. Super-resolution imaging of Sec61β-Rho-tag labeled with TMR in living U2 OS cells.**
The super-resolution movie was reconstructed from high-density SMLM data analyzed with the DeepSTORM 2D neural network. Scale bar: 2 µm.

### Supplementary Figures

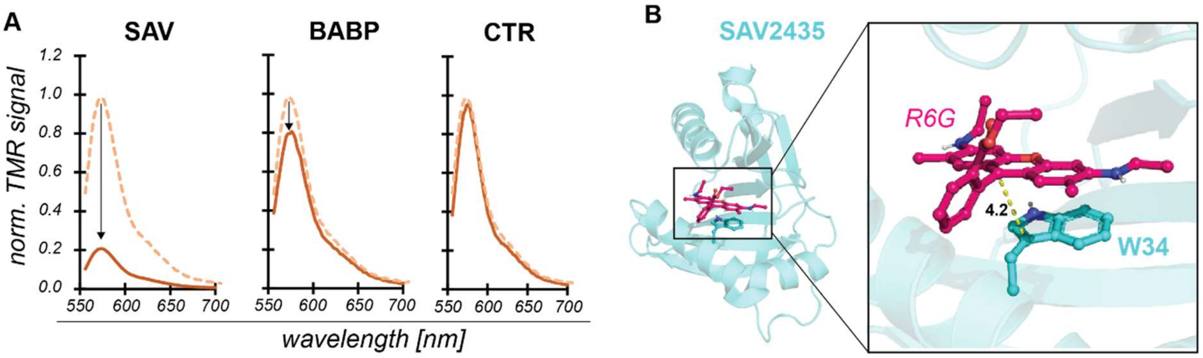

**Figure S1. The multidrug resistant protein SAV2435 quenches the fluorescence of bound dyes.**

A) Normalized fluorescence emission spectra of TMR (20 nM) in presence (straight line) or absence (dashed line) of the three initial Rho-tag candidates (20 µM). Black arrow indicates fluorescence decrease $F_I/F_{I0}$ upon protein binding.

B) Crystal structure and active site of the SAV2435 / R6G complex (PDB-ID 5KAW, 1.8 Å resolution). The tertiary structure is represented as cyan cartoon. The ligands and W34 residue are represented as colored sticks. Distances in Å.

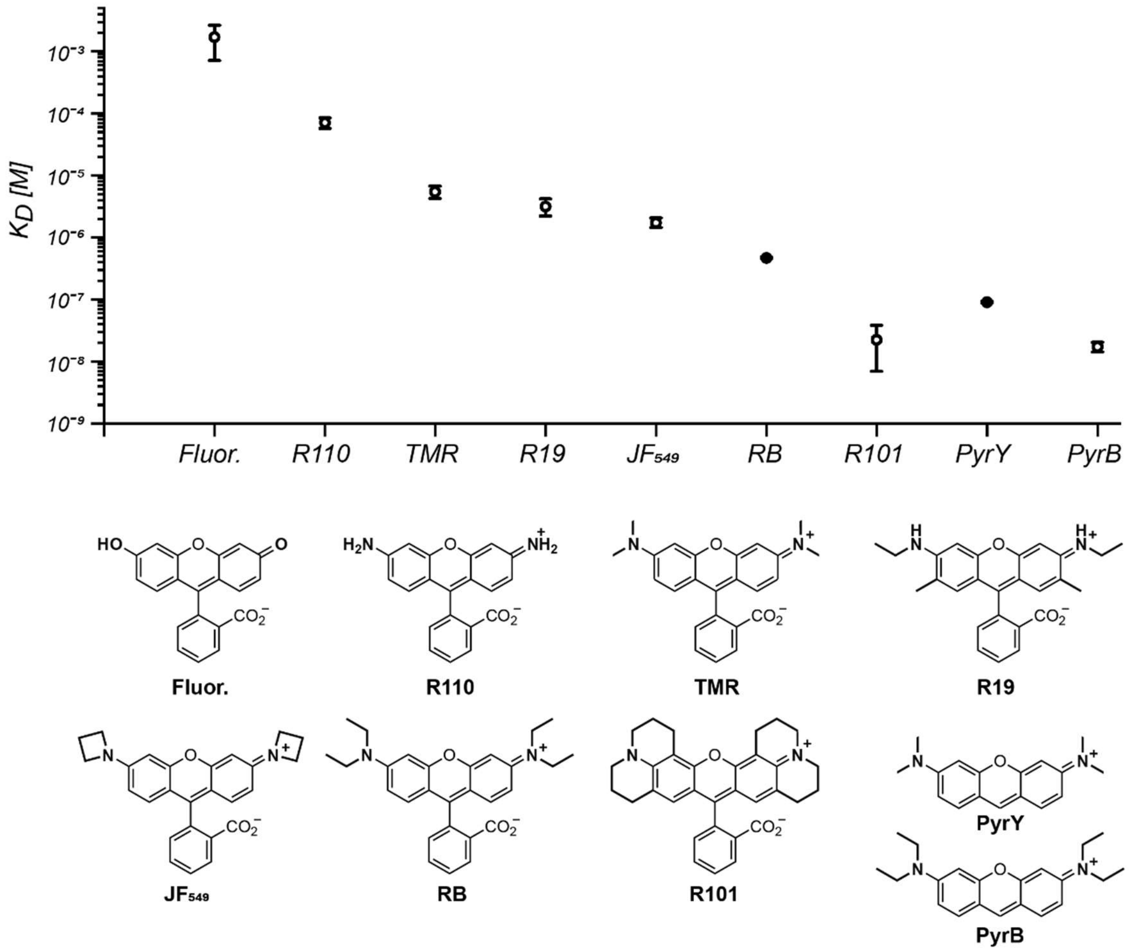

**Figure S2. Characterization of the dye-preference of CTR107.**

Binding affinity of CTR107 to a series of standard dyes. The fluorescence polarization of 20 nM dye (in buffer with 0.1% BSA) was recorded in presence of serial dilutions of CTR107 (0 – 200 µM). The $K_D$ was derived as the protein concentration at half-maximal FP. Data was fitted with Equation (2). Average data and standard deviation from at least three independent measurements (N≥3). Chemical structures of used dyes.

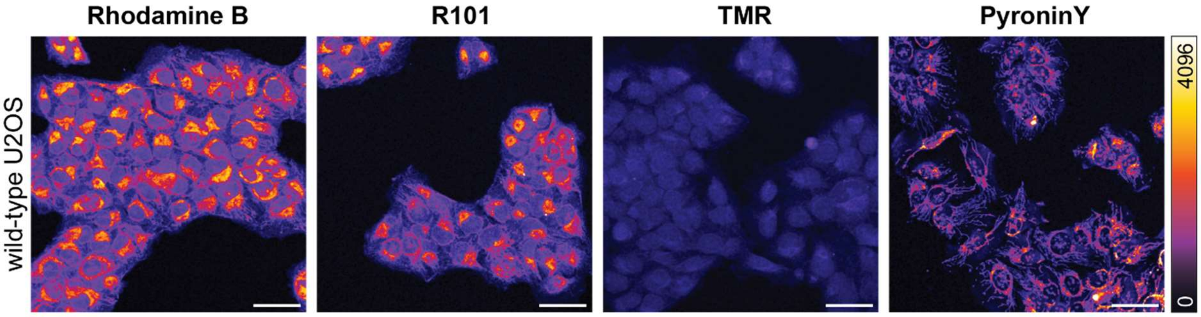

**Figure S3. Background signal of different rhodamine probes on U2OS cells.**

Live-cell confocal fluorescence of 200 nM of the indicated dyes on wild-type U2OS cells. Max. projections. Scale bars: 50 µm. Pixels colored according to reference bar (‘fire’ LUT).

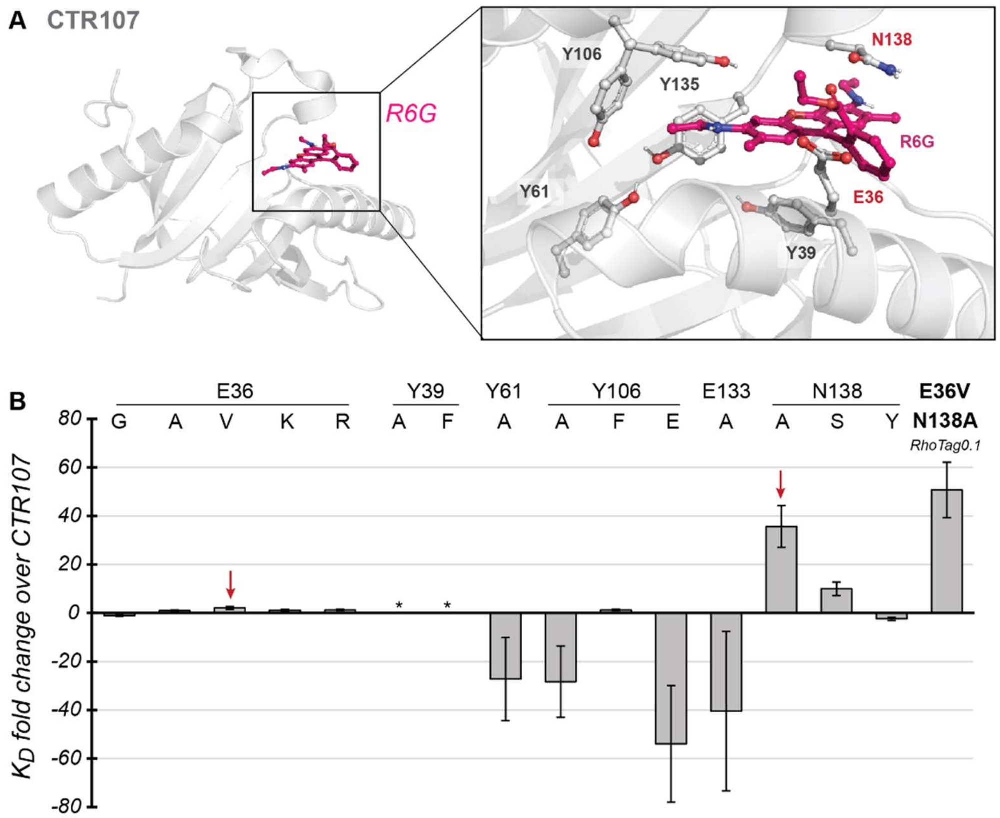

**Figure S4. Structure-aided side-directed mutagenesis screen for the generation of Rho-tag0.1.**

A) Crystal structure of CTR107 in complex with Rhodamine 6G (PDB-ID 5KAX, 2.0 Å resolution). Tertiary structures are represented as cartoons. Magnification of the fluorophore binding site and active site residues. Ligand and residues targeted by site-directed mutagenesis (except E133) as represented as sticks.

B) Fold-change in dissociation constant ($K_D$) to TMR (2 nM) for CTR107 point mutants (reference: 5.2 ± 0.7 µM). The $K_D$ was determined *via* FP assay. Average data and standard deviation from >3 replicates, error calculated using error propagation. E36V and N138A mutants (red arrows) were combined (Rho-tag0.1) yielding a ~48-fold improvement in TMR binding ($K_D$ 108 ± 29 nM).

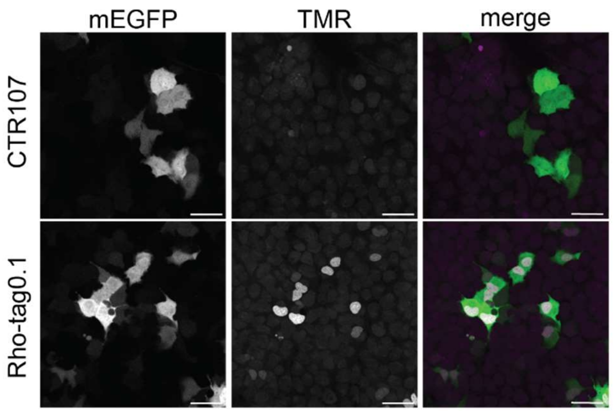

**Figure S5. Live-cell confocal imaging of CTR107 and Rho-tag0.1.**

U2OS cells expressing H2B-CTR107 (top) or -Rho-tag0.1-T2A-mEGFP (bottom) labeled with 200 nM TMR. Co-translational expression of mEGFP was used to identify Rho-tag expressing cells. Max. projections. Scale bars: 50 µm.

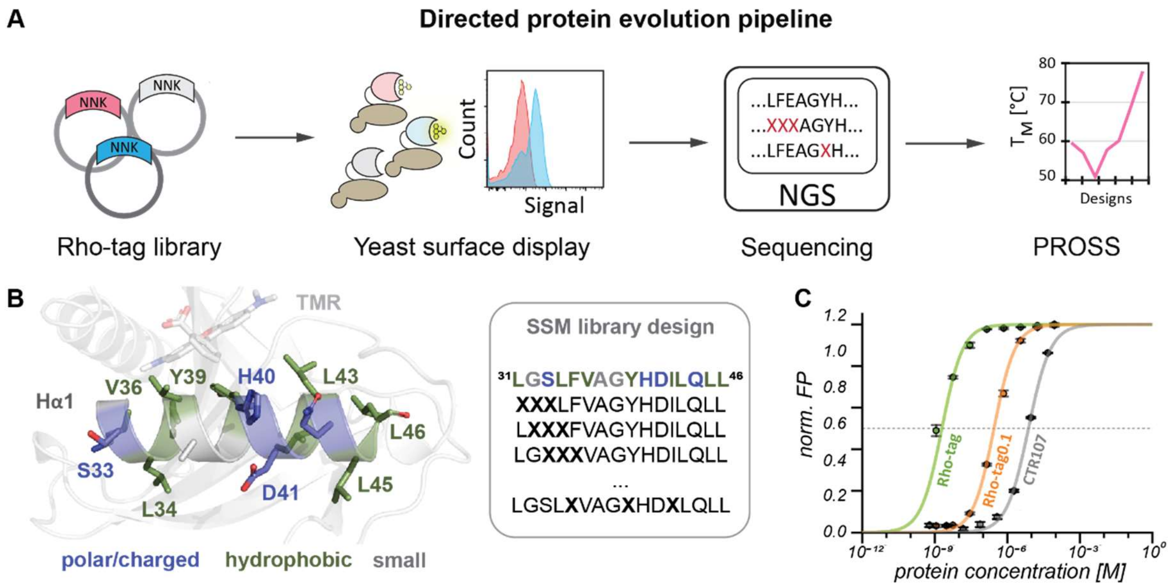

**Figure S6. High-throughput engineering of Rho-tag.**

A) Directed protein evolution strategy. Rho-tag0.1 randomized DNA libraries (sDMSL or SSML) were displayed on yeast surface and labeled with a fluorophore. Yeast cells were sorted and enriched for five consecutive rounds at decreasing dye concentrations. The selected variants were analyzed by next-generation sequencing (NGS) and decreased protein stability (melting temperature – $T_M$) was offset by using the Protein Repair One-Stop Shop (PROSS).[2]

B) Residues selected for randomization on *αH* and exemplary library design strategy. Crystal structure of CTR107$^{N138A}$ (PDB ID 9RTL), E36V mutation was inserted using Pymol mutagenesis function for representation purposes. X – NNK codon.

C) Fluorescence polarization titrations of Rho-tag candidates during development. The FP of TMR (0.5 nM) at varying protein concentrations was measured in presence of 0.1% BSA. Data was fitted with Equation (2) and normalized to the max./min. values of fitted curves. Representative curves of average data and standard deviation from technical triplicates.

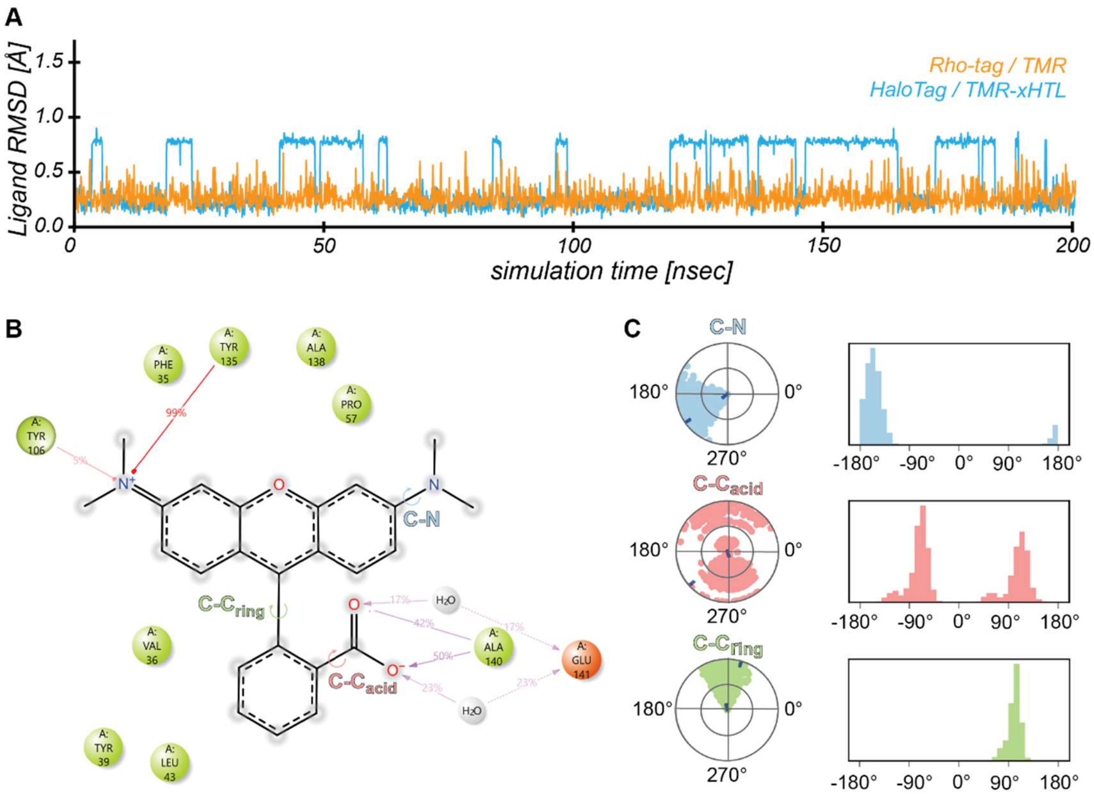

**Figure S7. Molecular dynamics simulation of the TMR protein interaction.**

A) Ligand RMSD over simulation time (200 msec) from MD simulation of the Rho-tag / TMR interaction (orange, PDB ID, 9RTM) compared to TMR-xHTL binding to HaloTag7 (blue, PDB ID 7ZIY). RMSD only calculated to the TMR moiety (21 atoms).

B) Ligand / protein interaction map. Interactions which appear for 5% of the simulation time displayed. Green ball – hydrophobic residue. Orange ball – charged residue. Grey ball – water molecule. Red line: π-cation interactions.

C) Ligand torsion profile. Radial plot and histogram representation of the conformational evolution around rotatable bonds as indicated in B (C-N, C-$C_{acid}$ and $CC_{ring}$).

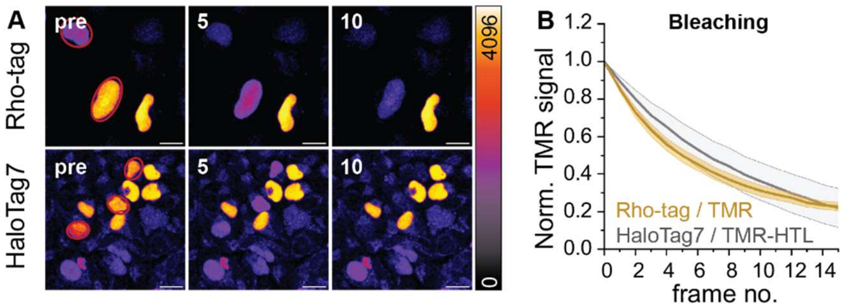

**Figure S8. Live-cell confocal bleaching comparison of Rho-tag and HaloTag.**

A) Live U2OS cells expressing Rho-tag- or HaloTag7-$P_{30}$-mEGFP-$NLS_3$ labeled with TMR or TMR-HTL, respectively (50 nM). Red ROI marks nuclei which were bleached with 100% laser power. Frame number displayed in upper left corner. Scale bars: 10 µm.

B) Quantification from data from A. Background corrected nuclear intensity was traced throughout bleaching process (frame number) and normalized to the first frame. N≥25 nuclei from 3 image series quantified.

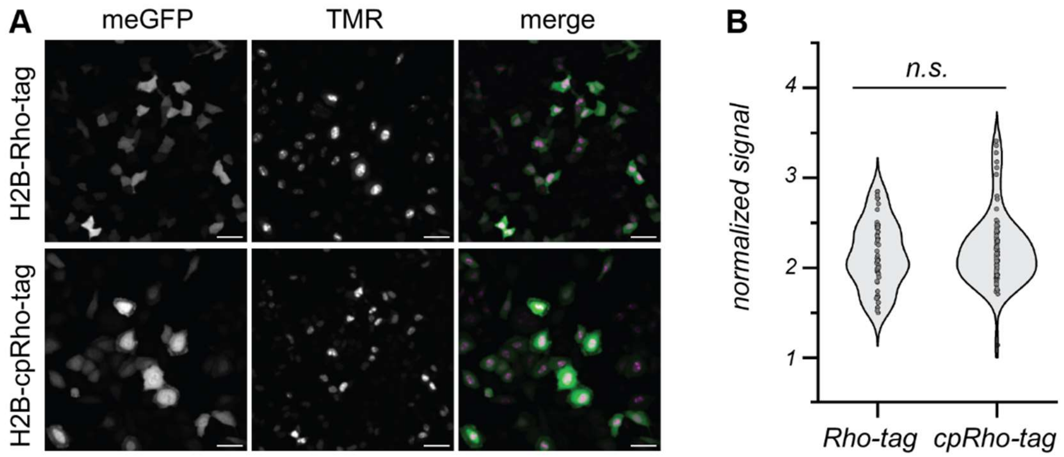

**Figure S9. Cellular signal comparison of Rho-tag and cpRho-tag.**

A) Cellular brightness comparison of live HeLa cells transiently expressing H2B-Rho-tag or -cpRho-tag labeled with TMR (50 nM). Co-translational mEGFP expression was used for signal normalization. Max. projections. Scale bars: 50 µm.

B) Cellular intensity comparison between Rho-tag and cpRho-tag labeling with TMR in live-cells. >60 nuclei were quantified from 3 images, global background subtracted and divided by the individual mEGFP signal. Significance was calculated using two-sided t-tests including the Welch correction. n.s.: $p \geq 0.05$, *: $p < 0.05$.

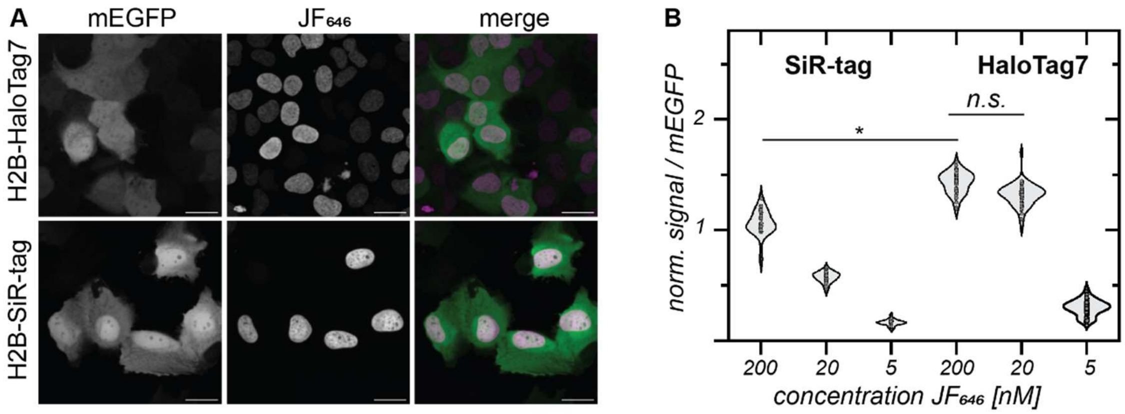

**Figure S10. Cellular signal comparison of SiR-tag and cpRho-tag.**

A) Cellular brightness comparison of live U2OS cells expressing H2B-SiR-tag or -HaloTag7 labeled with $JF_{646}$ (200 nM). Co-translational mEGFP expression was used for signal normalization. Max. projections. Scale bars: 50 µm.

B) Cellular intensity comparison between SiR-tag and HaloTag7 labeling with $JF_{646}$ in live-cells. >60 nuclei were quantified from 3 images, global background subtracted and divided by the individual mEGFP signal. Significance was calculated using two-sided t-tests including the Welch correction. n.s.: $p \geq 0.05$, *: $p < 0.05$.

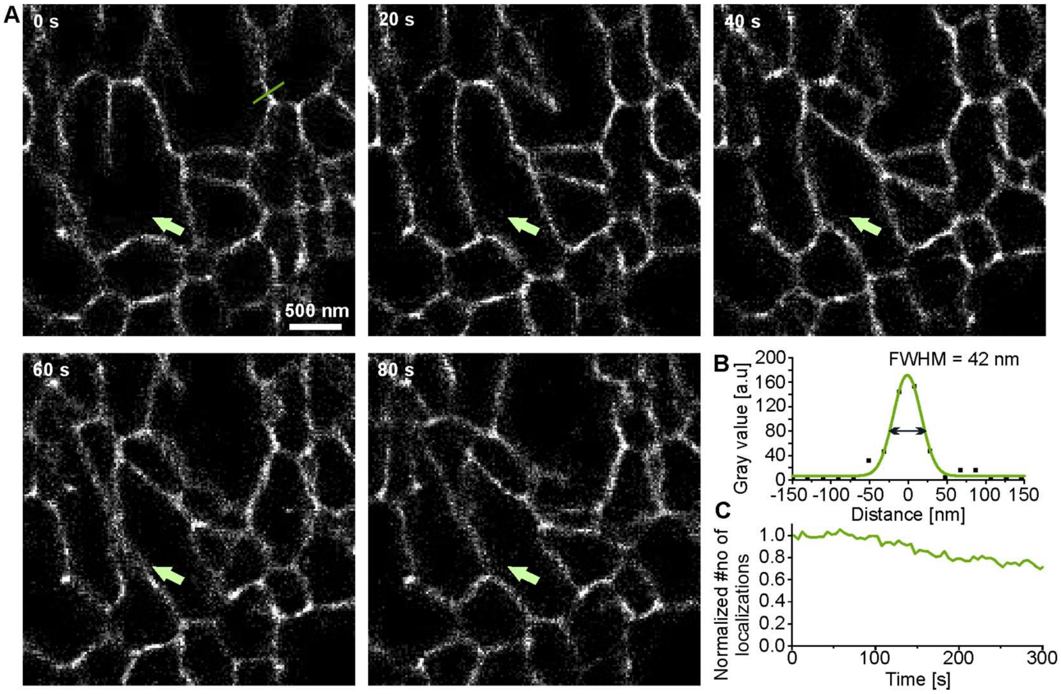

**Figure S11. Live-cell single-molecule SMLM imaging of Sec61β-Rho-tag labeled with TMR**

A) Super-resolution images of Sec61β-Rho-tag at different time points reconstructed from high-density single-molecule SMLM data. Analysis was performed with the neural network DeepSTORM 2D.[46]

B) Intensity profile of an ER tubule marked in A (green line) and the full-width at half maximum (FWHM).

C) The number of localizations over the acquisition time.

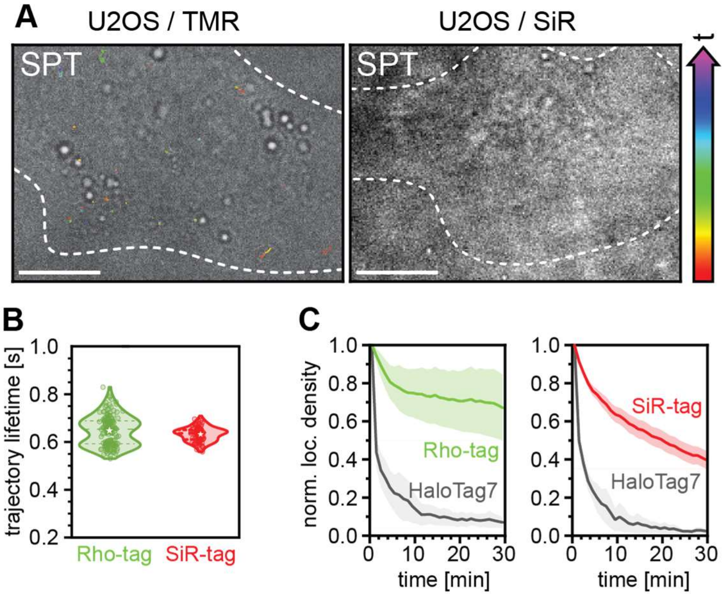

**Figure S 12. Live-cell single-particle tracking control experiments.**

A) Single-particle tracking (SPT) in living U2OS cells. Single-molecule trajectories recorded of TMR (left) or SiR on wild-type U2OS cells not expressing any Rho-tag / SiR-tag. Trajectories are color-coded by their time of occurrence as indicated by the colored arrow with early within a measurement appearing trajectories denoted in red and late appearing trajectories colored in purple. Scale bar: 10 µm.

B) Mean trajectory lifetime per cell for Rho-tag (0.648 ± 0.008 s) and SiR-tag (0.635 ± 0.008 s) and at a laser power of 0.5 kW/cm². Dotted lines in violin plots indicate the first and third quartiles, dashed lines the median, and stars the mean.

C) Number of localizations per area over time binned into 1 min intervals, normalized to the first interval and averaged over N = 5 cells. Comparison between Rho-tag / TMR and HaloTag7 / TMR-HTL (covalent) as well as SiR-tag / SiR and HaloTag7 / SiR-HTL (covalent).

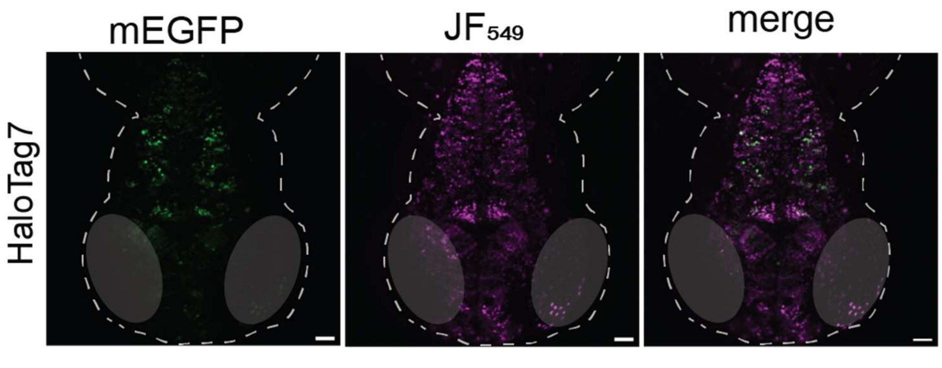

**Figure S 13. Full labeling of HaloTag in live zebrafish larvae.**

Live confocal imaging of zebrafish embryo mosaically expressing neuronal HaloTag7-$P_{30}$-meGFP-$NLS_3$ after overnight labelling, free-swimming in 10 µM $JF_{549}$-HTL. Scale bars: 50 µm.

### Supplementary Tables

**Supplementary Table 1 | Predicted physicochemical properties of rhodamine dyes and their HTL.**

| Molecule | Formula | MW [g/mol] | TPSA [Å²] | WLOGP | GI abs. | BBB | Pgp | log Kp [cm/s] | Ro5 Violat. |
|---|---|---|---|---|---|---|---|---|---|
| TMR-HTL | $C_{35}H_{42}ClN_3O_6$ | 636.2 | 107.1 | 4.2 | High | No | Yes | -9.60 | 1 |
| SiR-HTL$^{close}$ | $C_{37}H_{48}ClN_3O_5Si$ | 678.3 | 80.3 | 4.9 | High | No | Yes | -5.62 | 1 |
| TMR | $C_{24}H_{22}N_2O_3$ | 386.4 | 59.5 | 2.7 | High | Yes | Yes | -8.79 | 0 |
| SiR$^{close}$ | $C_{26}H_{28}N_2O_2Si$ | 428.6 | 32.78 | 3.3 | High | Yes | Yes | -4.81 | 1 |

*Data was generated with SwissADME[8]. MW – molecular weight. TPSA - total polar surface area. WLogP - predicted octanol/water partition coefficient. GI abs. – predicted passive gastrointestinal absorption (high/low). BBB – predicted probability for BBB permeation (Yes/No). Pgp - predicted as actively effluxed by P-gp (Yes/No). log Kp - skin permeability coefficient. Ro5 Violat. – number of violations to Lipinski's rule-of-five (0 -5).*

**Supplementary Table 2 | Predicted PROSS mutations for Rho-tag and melting temperature.**

| Design | Mutations | # | $T_M$ [°C] |
|---|---|---|---|
| **1** | V17Q, F62Y, G107D, S152V | 4 | - |
| **2** | Design 1 + S65D, E70D, V87I | 7 | 57.8 |
| **3** | Design 2 + Q25R, A66M, F125Y, D126E | 11 | - |
| **4** | Design 3 + K51V, G67E, D121K | 14 | 60.1 |
| **5** | Design 4 + S33E, G48E, S54A | 17 | - |
| **6** | Design 5 + A15E, A37K, T89V | 20 | 68.8 |
| **7** | Design 6 + L91E | 21 | - |
| **8** | Design 7 K37R + S29D, S52Q, G63N, G79P, A111K, R150Q | 27 | 77.8 |
| **9** | Design 8 G63H, K111R, R121K + Q6T, L45Y, S84K, S94A, D122A, D145E, Q146E | 34 | - |

*PROSS designs have a stacking number of mutations. S100C and A143P were excluded by manual inspection. $T_M$ was measured by nanoDSF. Reference: $T_M$ Rho-tag0.1 50.9 °C.*

**Supplementary Table 3 Computational designs of cplinker sequences for cpRho-tag**

| ID | pLDDT | seq | ID | pLDDT | seq |
|---|---|---|---|---|---|
| 228* | 96.00 | LGEVSGGEP | 43* | 94.51 | LSGPVTGSTP |
| 144 | 95.96 | LETADGEP | 120 | 94.40 | LETSGSGP |
| 47 | 95.93 | LGEVTGGEP | 24 | 94.26 | LESSRGGR |
| 173 | 95.93 | LGEVTGGEP | 83 | 94.21 | LETADGQP |
| 46 | 95.81 | LGEVTGGRP | 67 | 94.18 | TEGAPEVLESRP |
| **153*** | **95.71** | **LKNAGPETETLESQP** | 7 | 94.18 | TEGAPEVLESRP |
| 146* | 95.53 | LKESSDAPETLESRP | 142 | 94.16 | LESSRGGP |
| 64 | 95.25 | LKNDGPAETLESRP | 23 | 94.16 | LESSRGGP |
| 16 | 94.82 | LKGSTWNGRP | 29 | 94.16 | LESSRGGP |
| 218* | 94.78 | LDDEITGSQP | 188 | 94.00 | LGEVSGSQP |
| 237* | 94.72 | LETSGSQP | 94 | 93.85 | LETAPGGP |
| 27 | 94.59 | LETSRGGP | 12 | 93.85 | GVEGGEELESRP |

*Top24 sequences ranked by their pLDDT from 240 backbone generated using RFdiffusion, ProteinMPNN and ColabFold. * tested in vitro, sequence outperforming the others highlighted in bold.*

**Supplementary Table 4. SSM libraries design for the engineering of *Hα1.***

| # | design | # | design |
|---|---|---|---|
| 1 | $^{31}$**XXX**LFVAGYHDILQLL$^{46}$ | 11 | $^{31}$LGSLFVAG**XXX**DILQLL$^{46}$ |
| 2 | $^{31}$L**XXX**FVAGYHDILQLL$^{46}$ | 12 | $^{31}$LGSLFVAGYHD**XXX**LL$^{46}$ |
| 3 | $^{31}$LGSLF**XXX**YHDILQLL$^{46}$ | 13 | $^{31}$LGSLFVAGYHDI**XXX**L$^{46}$ |
| 4 | $^{31}$LGSLFV**XXX**HDILQLL$^{46}$ | 14 | $^{31}$LGSLFVAGYHDIL**XXX**$^{46}$ |
| 5 | $^{31}$LGSLFVAGY**XXX**LQLL$^{46}$ | 15 | $^{31}$LG**X**LF**X**AGY**X**DILQLL$^{46}$ |
| 6 | $^{31}$LGSLFVAGYH**XXX**QLL$^{46}$ | 16 | $^{31}$LGSL**X**VAG**X**HDI**X**QLL$^{46}$ |
| 7 | $^{31}$LG**XXX**VAGYHDILQLL$^{46}$ | 17 | $^{31}$LGSL**XX**AG**X**HDILQLL$^{46}$ |
| 8 | $^{31}$LGS**XXX**AGYHDILQLL$^{46}$ | 18 | $^{31}$LGSL**X**VAG**XX**DILQLL$^{46}$ |
| 9 | $^{31}$LGSL**XXX**GYHDILQLL$^{46}$ | 19 | $^{31}$**X**GSL**XX**AGYHDILQLL$^{46}$ |
| 10 | $^{31}$LGSLFVA**XXX**DILQLL$^{46}$ | 20 | $^{31}$LGSL**X**VAG**X**HD**X**LQLL$^{46}$ |

X – degenerated NNK codon. Libraries were transformed to EBY100 with at least 2-times coverage over the theoretical library size of $9.3 \times 10^3$.

**Supplementary Table 5| SSM libraries design for the engineering of *loop2.***

| # | design | # | design |
|---|---|---|---|
| 1 | [$^{133}$**XXXX**DAPAETAPDQLRT$^{149}$] | 14 | [$^{133}$EIYLDAPAETAPD**XXXX**$^{149}$] |
| 2 | [$^{133}$E**XXXX**APAETAPDQLRT$^{149}$] | 15 | [$^{133}$**X**I**X**LD**XX**AETAPDQLRT$^{149}$] |
| 3 | [$^{133}$EI**XXXX**PAETAPDQLRT$^{149}$] | 16 | [$^{133}$E**X**Y**X**D**X**P**X**ETAPDQLRT$^{149}$ |
| 4 | [$^{133}$EIY**XXXX**AETAPDQLRT$^{149}$] | 17 | [$^{133}$EI**X**L**X**APA**XX**APDQLRT$^{149}$] |
| 5 | [$^{133}$EIYL**XXXX**ETAPDQLRT$^{149}$] | 18 | [$^{133}$EIY**X**DA**XX**E**X**APDQLRT$^{149}$] |
| 6 | [$^{133}$EIYLD**XXXX**TAPDQLRT$^{149}$] | 19 | [$^{133}$EIYLD**X**P**X**E**XX**PDQLRT$^{149}$] |
| 7 | [$^{133}$EIYLDA**XXXX**APDQLRT$^{149}$] | 20 | [$^{133}$EIYLDA**X**A**X**T**XX**DQLRT$^{149}$] |
| 8 | [$^{133}$EIYLDAP**XXXX**PDQLRT$^{149}$] | 21 | [$^{133}$EIYLDAPA**XX**AP**XX**LRT$^{149}$] |
| 9 | [$^{133}$EIYLDAPA**XXXX**DQLRT$^{149}$] | 22 | [$^{133}$EIYLDAPA**X**T**XX**DQ**X**RT$^{149}$] |
| 10 | [$^{133}$EIYLDAPAE**XXXX**QLRT$^{149}$] | 23 | [$^{133}$EIYLDAPAE**X**AP**XX**L**X**T$^{149}$] |
| 11 | [$^{133}$EIYLDAPAET**XXXX**LRT$^{149}$] | 24 | [$^{133}$EIYLDAPAET**X**P**X**Q**X**R**X**$^{149}$] |
| 12 | [$^{133}$EIYLDAPAETA**XXXX**RT$^{149}$] | 25 | [$^{133}$EIYLDAPAE**XX**PDQL**XX**$^{149}$] |
| 13 | [$^{133}$EIYLDAPAETAP**XXXX**T$^{149}$] | | |

X – degenerated NNK codon. Libraries were transformed to EBY100 with at least 2-times coverage over the theoretical library size of 2.0 x $10^5$.

**Supplementary Table 6. SSM libraries design for the engineering of *β*.**

| # | design | # | design |
|---|---|---|---|
| 1 | $^{51}$**XXXX**GPPFARYYGFDMETFDVEFGFPVE$^{79}$ | 14 | $^{51}$VSPSGPPFARYYG**XXXX**TFDVEFGFPVE$^{79}$ |
| 2 | $^{51}$V**XXXX**PPFARYYGFDMETFDVEFGFPVE$^{79}$ | 15 | $^{51}$VSPSGPPFARYYGF**XXXX**FDVEFGFPVE$^{79}$ |
| 3 | $^{51}$VS**XXXX**PFARYYGFDMETFDVEFGFPVE$^{79}$ | 16 | $^{51}$VSPSGPPFARYYGFD**XXXX**DVEFGFPVE$^{79}$ |
| 4 | $^{51}$VSP**XXXX**FARYYGFDMETFDVEFGFPVE$^{79}$ | 17 | $^{51}$VSPSGPPFARYYGFDM**XXXX**VEFGFPVE$^{79}$ |
| 5 | $^{51}$VSPS**XXXX**ARYYGFDMETFDVEFGFPVE$^{79}$ | 18 | $^{51}$VSPSGPPFARYYGFDME**XXXX**EFGFPVE$^{79}$ |
| 6 | $^{51}$VSPSG**XXXX**RYYGFDMETFDVEFGFPVE$^{79}$ | 19 | $^{51}$VSPSGPPFARYYGFDMET**XXXX**FGFPVE$^{79}$ |
| 7 | $^{51}$VSPSGP**XXXX**YYGFDMETFDVEFGFPVE$^{79}$ | 20 | $^{51}$VSPSGPPFARYYGFDMETF**XXXX**GFPVE$^{79}$ |
| 8 | $^{51}$VSPSGPP**XXXX**YGFDMETFDVEFGFPVE$^{79}$ | 21 | $^{51}$VSPSGPPFARYYGFDMETFD**XXXX**FPVE$^{79}$ |
| 9 | $^{51}$VSPSGPPF**XXXX**GFDMETFDVEFGFPVE$^{79}$ | 22 | $^{51}$VSPSGPPFARYYGFDMETFDV**XXXX**PVE$^{79}$ |
| 10 | $^{51}$VSPSGPPFA**XXXX**FDMETFDVEFGFPVE$^{79}$ | 23 | $^{51}$VSPSGPPFARYYGFDMETFDVE**XXXX**VE$^{79}$ |
| 11 | $^{51}$VSPSGPPFAR**XXXX**DMETFDVEFGFPVE$^{79}$ | 24 | $^{51}$VSPSGPPFARYYGFDMETFDVEF**XXXX**E$^{79}$ |
| 12 | $^{51}$VSPSGPPFARY**XXXX**METFDVEFGFPVE$^{79}$ | 25 | $^{51}$VSPSGPPFARYYGFDMETFDVEFG**XXXX**$^{79}$ |
| 13 | $^{51}$VSPSGPPFARYY**XXXX**ETFDVEFGFPVE$^{79}$ | | |

X – degenerated NNK codon. Libraries were transformed to EBY100 with at least 2-times coverage over the theoretical library size of 2.0 x $10^5$.

**Supplementary Table 7 | Characterization of *putative SiR-tag variants.***

| | variant | sequence | $K_D$ [M] | $T_M$ [° C] |
|---|---|---|---|---|
| *loop2* | par | $^{133}$EIYLDAPAETAPDQLRT$^{149}$ | $6.5 \pm 2.1 \times 10^{-6}$ | 60.6 |
| | 1* | $^{133}$E**V**Y**M**D**Y**P**S**ETAPDQLRT$^{149}$ | $1.3 \times 10^{-6}$ | 60.2 |
| | 2 | $^{133}$EIYLD**SHY**ETAPDQLRT$^{149}$ | $9.1 \pm 3.5 \times 10^{-8}$ | 58.0 |
| | 3* | $^{133}$E**T**YLD**Y**P**R**ETAPDQLRT$^{149}$ | $8.7 \times 10^{-7}$ | 57.0 |
| | 4* | $^{133}$E**C**Y**W**D**W**P**T**ETAPDQLRT$^{149}$ | $3.7 \times 10^{-6}$ | 56.6 |
| | 5 | $^{133}$EIYLDA**YYWS**APDQLRT$^{149}$ | $3.6 \pm 0.6 \times 10^{-7}$ | 51.6 |
| | 6* | $^{133}$E**L**YLD**W**P**S**ETAPDQLRT$^{149}$ | $3.7 \times 10^{-7}$ | 53.4 |
| | 7* | $^{133}$EIYL**SFGV**ETAPDQLRT$^{149}$ | $6.3 \times 10^{-7}$ | 53.6 |
| | 8* | $^{133}$EIYL**SYFL**ETAPDQLRT$^{149}$ | $3.7 \times 10^{-6}$ | 55.1 |
| *β1* | 2 + PROSS8 | $^{51}$VSPSGPPFARYYGFDME$^{67}$ | $2.7 \pm 0.9 \times 10^{-7}$ | 75.6 |
| | 9* | $^{51}$VSPSGP**MYAL**YYGFDME$^{67}$ | $4.7 \times 10^{-8}$ | 70.5 |
| | 10* | $^{51}$VSPSGPPFARYY**NKNH**E$^{67}$ | $3.0 \times 10^{-8}$ | 71.0 |
| | 11* | $^{51}$VSPSGPPFARYYG**PRN**E$^{67}$ | $2.6 \times 10^{-7}$ | 56.7 |
| | 12* | $^{1}$VSPSGPPFARYYGF**S**M**R**$^{67}$ | $2.8\times 10^{-7}$ | 60.7 |
| | 13* | $^{51}$VSPSGPPFARYYG**YNPH**$^{67}$ | $3.8\times 10^{-7}$ | 72.5 |
| | 14* | $^{51}$VSPSGPPFARYYG**NSGK**$^{67}$ | $3.0\times 10^{-7}$ | 76.3 |
| | 9 + 10 (SiR-tag) | $^{51}$VSPSGP**MYAL**YY**NKNH**E$^{67}$ | $2.0 \pm 1.4 \times 10^{-8}$ | 71.2 |

*Binding affinity determined via fluorescence intensity. *data from single replicate. If not stated otherwise, average data and standard deviation from n ≥ 3 individual replicates. Thermostability ($T_M$) determined via nanoDSF measurements.*

**Supplementary Table 8. Data collection and refinement statistics for X-ray structures.**

| | **CTR107$^{N138A}$ / TMR** PDB ID 9RTL | **Rho-tag / TMR** PDB ID 9RTM |
|---|---|---|
| **Data collection** | | |
| Space group | $P6_5$ | $P2_12_12$ |
| Unit-cell parameters | | |
| *a, b, c* (Å) | 71.81, 71.81, 146.16 | 62.09, 62.35, 96.79 |
| $\alpha, \beta, \gamma$ (°) | 90.00, 90.00, 120.00 | 90.00, 90.00, 90.00 |
| Radiation source | PXII-X10SA, SLS | PXII-X10SA, SLS |
| Wavelength (Å) | 0.99999 | 1.00001 |
| Temperature (K) | 100 | 100 |
| Resolution range (Å) | 50-1.80 (1.90-1.80) | 50-2.10 (2.20-2.10) |
| No. of observed reflections | 410365 (63145) | 147000 (20041) |
| No. of unique reflections | 39388 (5878) | 22503 (2884) |
| Multiplicity | 10.4 (10.7) | 6.5 (6.9) |
| Completeness (%) | 99.9 (99.9) | 99.7 (100.0) |
| $R_{merge}$ (%) | 4.1 (39.7) | 5.9 (52.1) |
| $\langle I/\sigma(I) \rangle$ | 29.9 (6.0) | 16.7 (3.4) |
| $CC_{1/2}$ (%)$^{\#}$ | 100.0 (97.0) | 99.9 (90.5) |
| **Refinement** | | |
| Molecules per a.u. | 2 | 2 |
| No. of reflections | 39386 | 22503 |
| No. of reflections in test set | 1970 | 1132 |
| Resolution range (Å) | 38.35-1.80 | 44.00-2.10 |
| No. of non-hydrogen atoms | | |
| Total | 2554 | 2466 |
| Protein | 2343 | 2381 |
| Water | 123 | 27 |
| Ligand/ion | 88 | 58 |
| *R* (%) | 18.25 | 19.49 |
| $R_{free}$ (%) | 21.33 | 21.62 |
| RMS deviations from ideal | | |
| bonds (Å) | 0.013 | 0.009 |
| angles (°) | 1.337 | 0.913 |
| *B*-factors (Å$^2$) | | |
| Average | 36.29 | 46.87 |
| Protein | 36.21 | 47.23 |
| Water | 39.54 | 41.94 |
| Ligand/ion | 33.72 | 34.06 |
| Wilson B ( Å$^2$) | 32.16 | 42.13 |
| Ramachandran statistics (%) | | |
| favored regions | 98.7 | 96.4 |
| allowed regions | 1.3 | 3.6 |
| disallowed regions | 0 | 0 |
| Clashscore | 4.23 | 6.96 |

*Values in parentheses are for the highest resolution shell, $^{\#}$as implemented in XDS[51].*

**Chemical Structures**

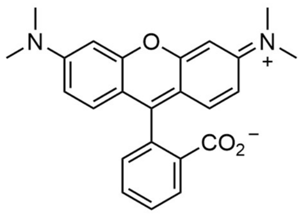

**TMR**

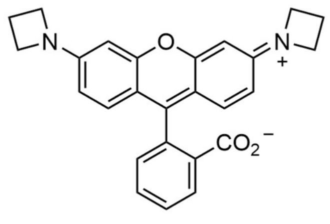

**$JF_{549}$**

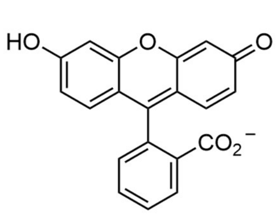

**Fluorescein**

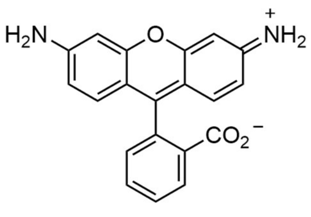

**R110**

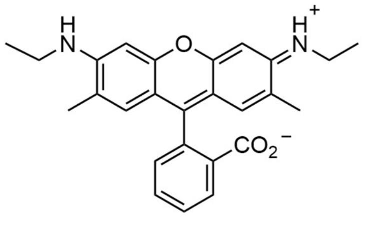

**R19**

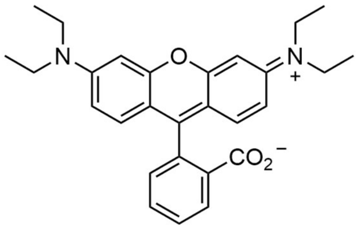

**Rhodamine B**

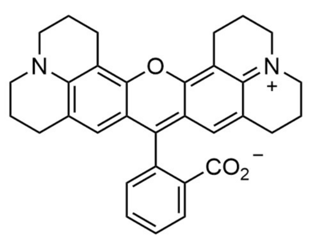

**R101**

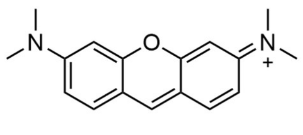

**Pyronine Y**

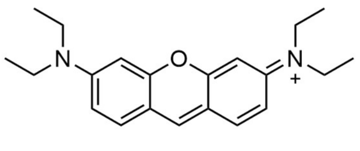

**Pyronine B**

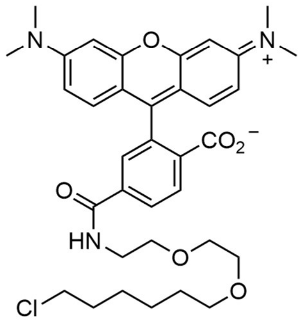

**TMR-6-HTL**

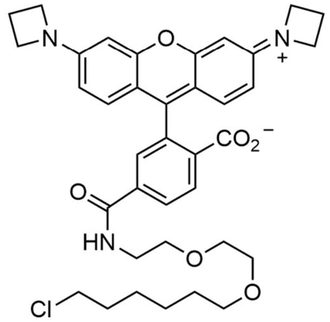

**$JF_{549}$-HTL**

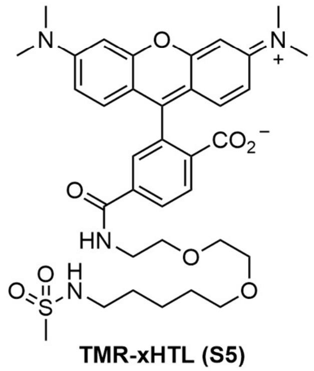

**TMR-xHTL (S5)**

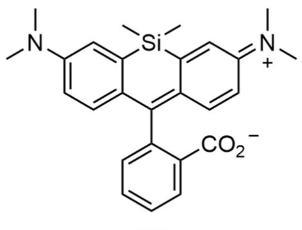

**SiR**

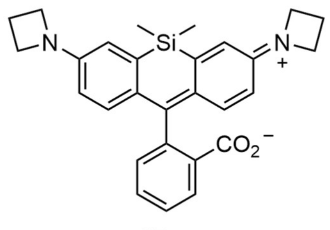

**$JF_{646}$**

**Figure S 14. Chemical structures dyes used throughout this study.** *Continues next page.*

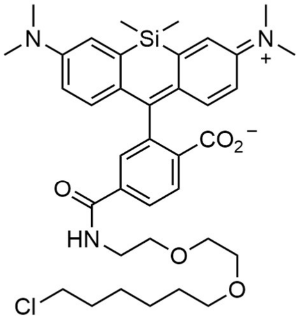

**SiR-HTL**

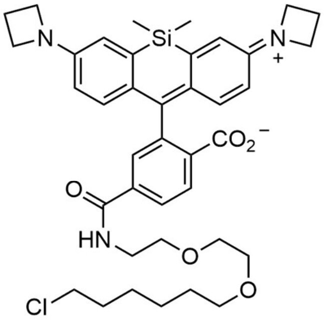

**JF$_{646}$-HTL**

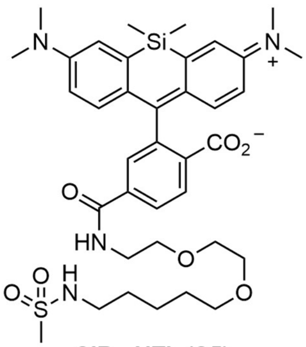

**SiR-xHTL (S5)**

**PROTEIN SEQUENCES**

*CTR107*
MDFECQFVCELKELAPVPALLIRTQTTMSELGSLFEAGYHDILQLLAGQGKSPSGPPFARYFGM
SAGTFEVEFGFPVEGGVEGSGRVVTGLTPSGKAASSLYIGPYGEIEAVYDALMKWVDDNGFDL
SGEAYEIYLDNPAETAPDQLRTRVSLMLHESE

*Rho-tag*
MDFECQFVCELKELAPQPALLIRTRTTMSELGNLFVAGYTDILQLLAGQGVSPSGPPFARYYGF
DMETFDVEFGFPVEGGVEGSGRIVTGLTPSGKAASSLYIGPYDEIEAVYDALMKWVKDNGYEL
SGEAYEIYLDAPAETAPDQLRTRVVLMLHES

*cpRho-tag (cplinker)*
ETFDVEFGFPVEPGVEGSGRIVVGETPSGKAASCLYIGPYDEIEKVYDALMKWVRDNGYELSG
EAYEIYLDAPAETPPDQLRTQVVLMLKNAGPETETLESQPALLIRTRTTMDELGNLFVAGYTDIL
QLLAEQGVQPAGPPFARYYNM

*RhoLuc (Rho-tag, NLuc)*
MDFECQFVCELKELAPQPALLIRTRTTMSELGNLFVAGYTDILQLLAGQGVSPSGPPFARYYGF
GSSGDQMGQIEKIFKVVYPVDDHHFKVILHYGTLVIDGVTPNMIDYFGRPYEGIAVFDGKKITVT
GTLWNGNKIIDERLINPDGSLLFRVTINGVTGWRLCERILAGGTGGSGGTGGSMVFTLEDFVGD
WRQTAGYNLDQVLEQGGVSSLFQNLGVSVTPIQRIVLSGENGLKIDIHVIIPYEGLGSETFDVEF
GFPVEGGVEGSGRIVTGLTPSGKAASSLYIGPYDEIEAVYDALMKWVKDNGYELSGEAYEIYLD
APAETAPDQLRTRVVLMLHES

*SiR-tag*
MDFECQFVCELKELEPQPALLIRTRTTMDELGELFVAGYHDILQLLAEQGVQPAGPMYALYYNK
NHETFDVEFGFPVEPGVEGSGRIVVGETPSGKAASCLYIGPYDEIEKVYDALMKWVRDNGYEL
SGEAYEIYLDSHYETPPDQLRTQVVLMLHESLE
